# Emergent catch bonds from slip-like contacts: cooperative force-driven proofreading at receptor–ligand interfaces

**DOI:** 10.64898/2026.09.15.751850

**Authors:** Juan Iribas, Miguel García-Sánchez, José Faro, Mario Castro

## Abstract

Mechanical forces modulate receptor-ligand kinetics, sometimes producing the so-called catch bond: a tensile force strengthens the bond, whereas in most cases engagement time is reduced (a slip bond). Multiple phenomenological models reproduce catch-slip behavior between receptors and ligands, primarily by positing how the reaction’s free-energy landscape changes under applied forces. In this paper, we show that distinct microscopic mechanisms often collapse into mathematically equivalent multi-exponential forms. This operational equivalence makes them difficult to distinguish (or even falsify) based solely on bond survival curves. We expose this mathematical equivalence and the validity regimes of these reductions and introduce *c*ooperative *f*orce-driven *p*roof*r*eading (CFPR), a network-level mechanism where pulling geometry enables contact-zone reorganization and additional nucleation of residue binding contributions, generating emergent catch-like responses from individually slip-like residue binding contributions. Our model can incorporate data from molecular dynamics simulations to predict, in advance, whether catch-slip will be present in a system and under what conditions, making it a predictive tool that complements experimental data.

**SIGNIFICANCE:** Living cells constantly pull on the bonds between receptors and their molecular partners. Counterintuitively, some bonds become longer-lived when pulled (catch bonds), allowing immune cells and adhering cells sense and respond to mechanical force. Many competing theories reproduce the same lifetime measurements, so which mechanism actually operates is often unknown. We show mathematically why these theories are frequently indistinguishable from bond-lifetime data alone, identify where widely used approximations break down, and propose a new mechanism in which force exposes new molecular contacts that reinforce binding. Because it directly links to molecular dynamics simulations, our framework can predict in advance when catch bonds form, guiding experiments and the design of force-sensitive molecules.

## 1 INTRODUCTION

For most receptor-ligand interactions, tensile force accelerates dissociation: a larger force corresponds to a shorter bond lifetime, a mechanism explained in terms of free-energy landscapes and mechanical work by Bell (1). However, another bond type violates this intuition by initially decreasing the effective *off* -rate *k* ^−^ over a finite force range (catch bonds), in contrast with the former (slip bonds). Dembo et al. predicted catch bonds within a Bell-type framework (2), and experiments later observed them in bacterial adhesion (FimH–mannose) (3) and in selectin-mediated leukocyte adhesion (4). The response is usually biphasic, with a catch-to-slip transition at higher forces (catch-slip bonds), as shown schematically in Fig. 1b. More recently, catch-slip behavior was observed to also occur in integrins, motor and cytoskeletal proteins, and T cell receptors (TCRs) (5, 6). These observations challenge the view that force merely lowers unbinding barriers — mechanical load can reorganize molecular interactions to stabilize binding over specific force ranges.

**Figure 1.**
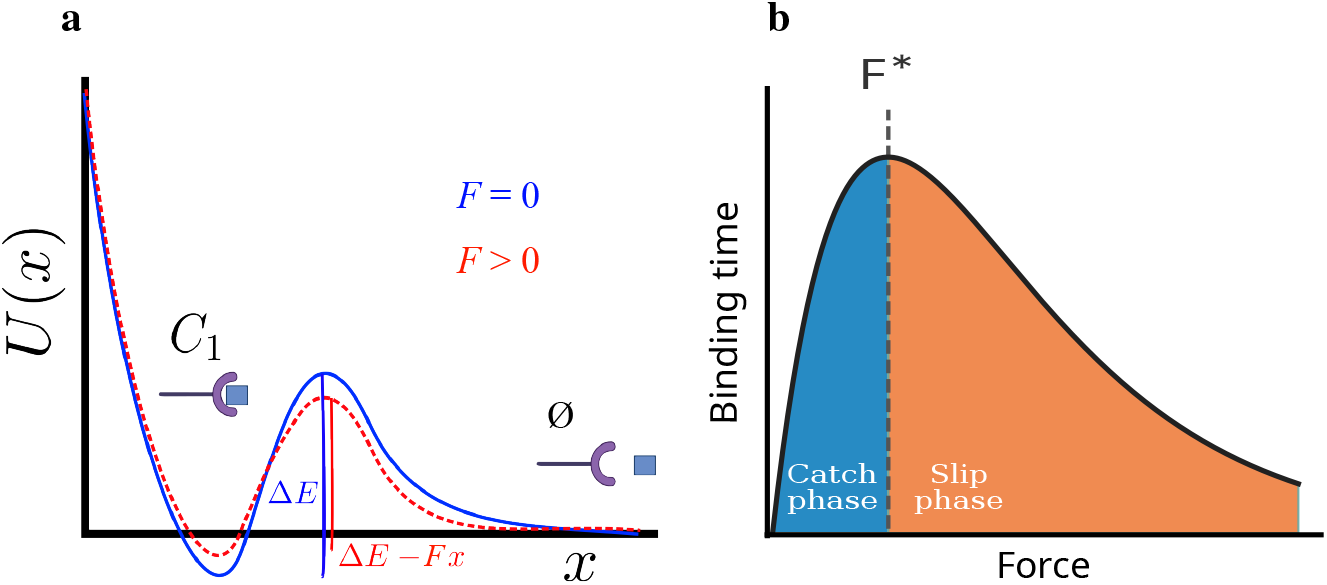
One-dimensional energy landscape and catch-bond schematic. **a)** Projecting the receptor–ligand landscape onto a reaction coordinate *x* yields a one-dimensional potential *U*_0_ (*x*) with a bound minimum separated by a barrier Δ*E* from the unbound state (blue line). An applied force tilts the landscape via *U* (*x*) = *U*_0_ (*x*) − *Fx*, lowering the effective barrier and accelerating dissociation (Bell model; red, dotted line). **b)** Schematic catch-bond force–lifetime curve. The mean bond lifetime first increases with force (catch phase, blue) before decreasing at higher forces (slip phase, orange), defining a characteristic peak force for bond stabilization.

Mechanical control of bond lifetimes is not merely a biophysical curiosity (6) but entails a well-recognized functional role. For instance, T lymphocytes form catch-slip bonds between TCR and peptide-MHC ligands on antigen-presenting cells (7, 8). These observations motivate the broader question of how force-dependent kinetics shape receptor discrimination in mechanically active cellular contexts, where bond lifetimes under load can control both information transmission and mechanical stability.

Multiple mechanisms explain catch-slip behavior, including two-pathway and two-state schemes, allosteric switching, deformation-controlled barriers, and sliding–rebinding (9–15). Although these models reproduce similar lifetime–force curves, they appear in different formalisms tailored to specific systems. This creates two problems. First, it obscures which assumptions are genuinely distinct and which reparametrize the same kinetics. Second, it renders inference *non-identifiable*: distinct mechanisms produce indistinguishable observables (16, 17). Force–lifetime data alone cannot discriminate between competing explanations.

In this paper, we show that force-lifetime data alone cannot independently constrain the key timescales underlying Kramers-type reductions (*e*.*g*., intrawell relaxation versus escape), because distinct parameterizations produce indistinguishable survival curves and, consequently, it makes model discrimination almost unattainable as one can fine-tune the fitting parameters without a connection with the microscopic details. Therefore, based only on that data, it is not possible to clarify the potential role of catch bonds in systems where the experiments have not been carried out (18). This motivates both a unified description of existing catch-bond models and complementary mechanisms connecting mesoscale observables to molecular structure.

This work has three aims. First, we survey major classes of catch-slip models and quantify the conditions under which Kramers-type reductions remain valid. Second, we show that under those conditions, many landscape-based mechanisms collapse onto a common multi-exponential kinetic form, implying broad operational equivalence. Third, we introduce a new mechanism (and the corresponding mathematical formulation) named Cooperative Force-Driven Proofreading (CFPR), a network-level mechanism not reducible to Kramers type form, adapted from kinetic proofreading (19, 20), and that emphasizes the role of cooperativity between hydrogen bonds, salt bridges, hydrophobic contacts — each called a *residue binding contribution* (RBC). Our model not only departs from unobserved energy landscape-based descriptions but also provides a direct connection to molecular dynamics simulations, making it both an explanatory and a predictive tool for the existence of catch-slip behaviors.

## 2 BACKGROUND: FORCE-DEPENDENT KINETICS OF RECEPTOR–LIGAND BONDS

We separate translational diffusion encounter and rotational diffusion orientation from chemical binding (21). Receptor (*R*) and ligand (*L*) first form a short-lived encounter complex *RL*^*\**^, then a properly oriented complex *RL*, and finally a bound complex *C* that can signal. This coarse-grained description separates geometric constraints, molecular flexibility, and chemical specificity:

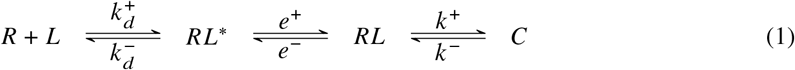

Here 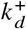 and 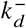 control formation and breakup of the encounter complex; *e*^+^, *e*^−^ govern orientation; and *k*^+^, *k* ^−^ control the transition to and from the bound state. Except *k*^+^ and *k* ^−^, all other rates depend on dimensionality (3D solution *vs* 2D membrane). Actually, the rate 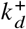 depends on reactant densities, while *k* ^−^ depends on mechanical load.

At a microscopic level, dissociation follows stochastic motion on a high-dimensional free-energy landscape. Typically, this dynamics is *projected* onto an effective reaction coordinate *x* along the dissociation pathway, giving a one-dimensional potential *U*_0_(*x*) (Fig. 1a). The Kramers zero-force escape rate is

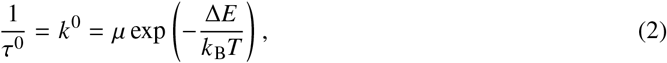

where Δ*E* is the energy barrier height, *µ* is a characteristic frequency (that depends on the local curvatures of the energy landscape at the minima), *k*_B_ is Boltzmann’s constant, and *T* is temperature (22). A required assumption is timescale separation: the intrawell relaxation time, *τ*_relax_, has to be fast compared with the mean escape time, *τ*_escape_, that is, *τ*_relax_ *≪ τ*_escape_. As the force rises or falls, this separation can break down (Section 3). Note that this view makes another strong assumption: it presupposes that the high-dimensional energy landscape is a smooth differentiable manifold with well-defined local minima and that the reaction coordinate *x* can be straightforwardly matched to the experimental distance between ligand and receptor (so the work performed by the force is proportional to *Fx*).

A tensile force *F* applied along the reaction coordinate tilts the landscape,

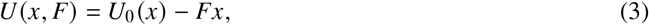

so the barrier is modified by the mechanical work done by the force. The effect of force is therefore encoded geometrically by projecting molecular deformation onto the reaction coordinate. If the dominant effect is a force-independent transition-state distance *x*, the classical Bell model follows (1, 23):

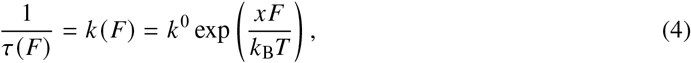

which predicts slip-bond behavior: increasing *F* increases *k* (*F*) and decreases the lifetime. Note that in Bell’s original formulation,

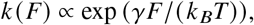

and later the coefficient *γ* was reinterpreted as the *reaction coordinate* and so *γF* as a clear-cut interpretation as the applied work.

In this single-pathway limit, the survival probability (or survival function) is mono-exponential,

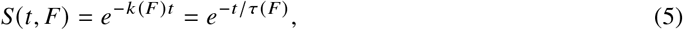

with a force-dependent time constant.

Real receptor–ligand systems access multiple microscopic pathways. A simple example is the two-state, two-pathway model (Fig. S3), described by the coupled reactions:

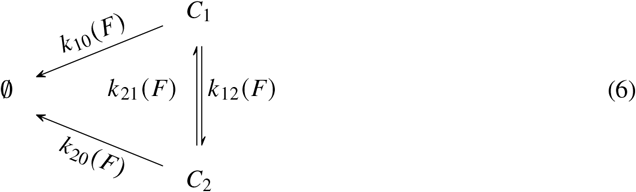

In this case, the survival probability is given by (see Supplementary Information, subsection S2.2 for details):

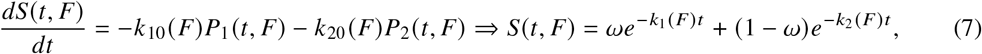

where *k*_1,2_ (*F*) are functions of the original rates *k*_10_ (*F*) , *k*_20_ (*F*) , *k*_21_ (*F*) and *k*_12_ (*F*) , and represent a fast and slow decay rate.

This landscape-based framework can be made as sophisticated as desired by choosing the shape and how the force deforms the landscape, but ultimately, under Kramer’s approximation, they all can be modeled by an underlying stochastic process that obeys a multidimensional master equation of the form

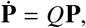

where **P** = (*P*_1_, … , *P*_*n*_) is the vector of probabilities of being in a specific state and *Q* the Markov generator of the process (24). From this master equation, it is straightforward to compute the survival function 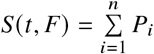 , which has a multi-exponential form:

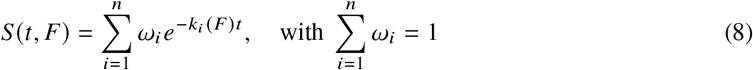

where the *k*_*i*_ (*F*) are the eigenvalues of *Q* and, biologically, are effective force-dependent rates. Catch-slip behavior emerges through (i) force-induced changes in the rates *k*_*i*_ (*F*), (ii) redistribution of weights *ω*_*i*_ among modes, or (iii) both. The simplest two-mode realization of Eq. (8) produces a biphasic survival curve whose force-dependent fast and slow phases are illustrated schematically in the Supplementary Material (Fig. S4).

In general, catch-slip behavior can arise from different microscopic or mesoscopic mechanisms. At the observable level, most datasets reduce to survival functions *S* (*t, F*), mean lifetimes *τ* (*F*) , or effective off-rates *k* (*F*) under prescribed loading. This creates two challenges. First, distinct physical pictures map onto the same *effective* kinetics (*e*.*g*., the same multi-exponential form in Eq. (8)). Second, mechanistically distinct models may be *non-identifiable* from force–lifetime data: different formulations of the underlying process produce indistinguishable predictions (16, 17). This is especially relevant when independent constraints on intrawell relaxation and escape timescales are scarce, so model fitting alone cannot be used to test the validity of the theory.

In the following section (and the Supplementary Material), we show that the main mechanistic models proposed to date can in fact be described by Eq. (8). Next, we introduce a new catch-bond model based on a cooperative, force-driven mechanism that does not rely on that equation for validation. Rather, this model can be falsified using independent molecular dynamics simulations and, in addition, can predict new systems in which the catch-bond effect can arise.

## 3 CATCH-SLIP MODELS: EQUIVALENCES AND VALIDITY REGIMES

The main catch-bond models are briefly described below. A detailed analysis, provided in Sec. S2 in the Supplementary Material, allows to extract their typical signatures and hence to detect identifiability issues in each of them. Table 1 lists them, summarizes their core physical mechanism, and provides key references.

**Table 1.** Summary of main catch-slip bond models. Representative references are listed; see also recent reviews (5, 6). More details in the Supplementary Material.

| Model class | Core physical idea | Refs. |
| --- | --- | --- |
| Two-pathway (TP) | Two dissociation pathways compete; force biases flux between a “catch” and a “slip” route. | (9, 10) |
| Two-state, two-pathway (TSTP) | Two bound conformations interconvert; each can dissociate via its own pathway. | (11) |
| Allosteric deformation (AD) | Force acts as an allosteric effector that stabilizes particular conformations by deformation energy. | (13, 14) |
| Bond-deformation (BD) | Force changes barrier height and bond energy (1D landscape). Force modifies local binding-pocket geometry, strengthening or weakening non-covalent contacts. | (12) |
| Sliding–rebinding (SR) | Force drives sliding along a structured interface, enabling repeated breakage/rebinding before full dissociation. | (15) |

### Two-pathway (TP)

This is an energy-landscape model that assumes a one-dimensional potential *U* (*x*) along the reaction coordinate *x*, with a minimum at *x* = 0 (bound step) and a barrier at *x*_*c*_ < 0 (catch) and another at *x*_*s*_ > 0 (slip). An applied force tilts the landscape, raising one barrier while lowering the other (Fig. S2). As shown in the Supplementary Material, section S2.6, the survival rate produced by this model is a single exponential. So, when a biphasic decay is observed, this model cannot even capture the experimental data (except in the accidental case where the two timescales are almost identical).

### Two-state, two-pathway (TSTP)

This model extends the two-pathway framework by resolving the bound state into two interconverting conformations, *C*_1_ and *C*_2_ (interconversion rates *k*_12_, *k*_21_), each dissociating through its own force-dependent Bell pathway with rates *k*_10_, *k*_20_ (see Supplementary Material, subsection S2.2). Such conformational multiplicity is well documented in receptor–ligand systems, from the TCR to antibodies and BCRs (25, 26). Solving the coupled master equations yields a genuinely *biphasic* survival probability,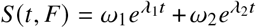 —the simplest two-mode realization of Eq. (8), with a fast decay from depletion of the short-lived state followed by a shallow tail set by the long-lived one. Catch behavior arises when increasing force shifts the statistical weight toward the more force-resistant conformation and/or lowers its off-rate, prolonging the mean lifetime over a range of forces. Because the two states differ only through their kinetic parameters, barrier heights, transition-state distances, and interconversion rates can trade off against one another; the model is therefore non-identifiable when *τ* (*F*) is the only output. In this model, the biphasic decay of the survival function is genuinely distinct from that of the TP model but identical to that of the other models below.

### Allosteric deformation (AD)

Building on the Monod–Wyman–Changeux picture (27), this model posits two receptor conformations —a tense, low-affinity state *R*_1_ and a relaxed, high-affinity state *R*_2_— that interconvert allosterically under load, with force biasing the receptor toward the relaxed state (see Supplementary Material, subsection S2.3). Its distinctive feature relative to the TSTP model is that conformational conversion and bond dissociation are governed by *separate* potentials, so the bound complex experiences a time-dependent effective off-rate *k* (*t, F*) = *k*_10_ (*F*) *Q*_1_ (*t, F*) + *k*_20_ *F*) *Q*_2_ (*t, F*) . Catch behavior emerges when force simultaneously stabilizes the high-affinity state and modulates the dissociation barriers, prolonging the lifetime before slip-like behavior sets in. Although its biophysical interpretation is distinct, the AD governing equations map exactly onto those of the TSTP model (subsection S2.6); it therefore generates identical bond survival statistics and shares the same identifiability problem when bond lifetime *vs* force is the only observable.

### Bond-deformation (BD)

This model is also based on the free-energy landscape framework, and assumes that force induces deformations of the binding pocket that strengthen or weaken the RBCs in the binding interphase. Although this model uses a one-dimensional potential with a single pathway (Fig. S1a), it includes in the Bell-like dissociation rate an additional force-dependent energy term, Δ*E*_*d*_ (*F*), to account for the change in the binding energy due to the binding-pocket deformation induced by the applied force (see details in the Supplementary Material, subsection S2.4). Similar to the AD model, but in contrast to other energy landscape models, this model postulates that both the maximum and the minimum of the potential well are lowered by the applied force, but by different mechanisms (the maximum by the Bell mechanism and the minimum by bond deformation) and at different rates. The deformation energy Δ*E*_*d*_ (*F*) can produce effective catch behavior if it dominates the Bell tilting term until a critical force *F*_0_ is attained, beyond which the Bell term dominates. For slow ramp forcing (force changing linearly with time), it produces a biphasic decay, but otherwise, using binding lifetime curves, it is indistinguishable from the TP model. Moreover, it has been shown that the AD model is a particular instance of the more general BD model (13). Hence, the BD model is also non-identifiable when bond lifetime *vs* force data is used as the only output.

### Sliding–rebinding (SR)

In the above models, reaction coordinates lack direct physical interpretation because they do not consider the receptor-ligand interface molecular structure. To address this issue, the SR model considers a detailed interface structural mechanism. Thus, the interface is represented by *N* pseudoatom pairs, each contributing with the same type of non-covalent bond. Under traction force, when all *N* pseudoatomic interactions dissociate, the complex can either fully dissociate or slide and form new interactions, this time at most *N* − 1 interactions. The model also considers rebinding of some pseudoatom pairs at the edge of the contact surface. This process is repeated until no further sliding is possible (only one pseudoatom pair remains) and rebinding does not occur. Thus, the receptor-ligand complex would slide, rebind, or fully dissociate. This model introduces multiple intermediate timescales and protocol dependence; effective reductions can mimic other multi-state models if intermediates are not resolved. Non-Bell-like effective rates may distinguish SR from TSTP. The effective rates of the reduced sliding–rebinding model (Appendix S6.4) involve ratios of force-dependent Bell rates (*k*_eff_ *∝ k*_−1_*k*_−2_/(*k*_+1_ + *k*_−2_)), producing concavity in semi-log plots over the crossover force range where *k*_+1_(*F*) *∼ k*_−2_(*F*), with scale *F*_cross_ *∼* (*k*_B_*T*/(*x*_+1_ +*x*_−2_)) ln 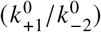. Both low- and high-force asymptotes are Bell-like but with different effective distances, making the intermediate curvature a diagnostic signature that allows distinguishing SR from TSTP and AD models when *F*_cross_ falls within the experimental force range.

At fixed force, the experimental survival curves of many of the above mechanistic models (TP, TSTP, AD, SR) can be well approximated by Eq. (8), either exactly or after timescale separation (Appendix S1). However, such a Kramers-type approximation in those catch-bond models is valid only if there is a proper timescale separation between fast intrawell relaxation (*τ*_relax_) and slow barrier escape (*τ*_escape_). The conditions under which this requirement holds can be determined as follows. For a one-dimensional potential with barrier height Δ*E* (*F*) = Δ*E*_0_ − *xF* and local curvature *ω* at the minimum, the relaxation time scales as *τ*_relax_ *∼* 1/*ω*, and the escape time as *τ*_escape_ *∼* exp[Δ*E* (*F*)/(*k*_*B*_*T*)]. Therefore, the validity criterion for this assumption (that is, *τ*_relax_ *≪ τ*_escape_) amounts to:

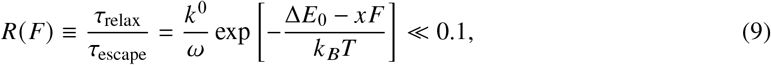

with *k*^0^ defined in Eq. (2).

Using representative parameters from the protein-ligand unbinding literature (Δ*E*_0_ ≈ 15*k*_*B*_*T* , *x* ≈ 0.5 nm, *ω* ≈ 10_12_ s^−1^ (23, 28)), the ratio *R* (*F*) remains well below 0.1 across the physiologically relevant force range, confirming that the timescale separation holds throughout 0–40 pN for typical receptor–ligand systems. Quantitative predictions in the high-force regime (beyond *∼*30 pN) should nonetheless be verified against full Fokker-Planck or molecular-dynamics treatment. For each of the models TP, TSTP, AD, and SR, we computed *R* (*F*) over the range 0–40 pN and confirmed that it remained well below 0.1 throughout this range for the stated parameters. Hence, in those models, the Kramers-type approximation is valid.

This lends additional support to the above mentioned identifiability problem, affecting similarly to all those models: in the absence of independent information about relaxation and escape timescales, *good fits* at the level of *τ* (*F*) (or the off-rate *k* (*F*)) does not guarantee unique physical interpretation of inferred parameters because barrier heights, transition-state distances, and interconversion kinetics can compensate one another (16, 17). Therefore, to discriminate between models, additional observables are required besides *τ* (*F*) , for instance: (i) deviations from mono-exponential survival at fixed force (revealing hidden states), (ii) protocol dependence (constant force versus ramp, pulling geometry, or cycling force regime), and (iii) independent structural or dynamical experiments. In the TCR context, pulling geometry is especially informative because it can alter alignment and contact formation at the interphase, motivating cooperative mechanisms that predict a slip-to-catch transition when pulling promotes additional bonding. This is addressed in the next section.

## 4 COOPERATIVE FORCE-DRIVEN PROOFREADING (CFPR)

In the previous section, we showed that landscape-based catch-bond mechanisms collapse onto similar multi-exponential kinetics under timescale separation, making inference of mechanisms from force–lifetime data fragile. Here we introduce a *network-level* mechanism at the protein-protein bond interphase, denoted *Cooperative Force-Driven Proofreading* (CFPR). Within the TCR–pMHC contact zone, the receptor–ligand bond comprises multiple individual non-covalent micro-contacts or RBCs — hydrogen bonds, salt bridges, hydrophobic contacts. Like Hopfield/Ninio kinetic proofreading (19, 20), our mechanism is also a non-equilibrium process, but instead of consuming ATP, it consumes mechanical work and dissipates entropy (see Eq. (24)).

### 4.1 Birth–death description

We consider that the TCR-pMHC bond interphase contains a small network of RBCs. We coarse-grain this network into a one-dimensional ladder of states labeled by the number of engaged RBCs, *n ∈* {0, 1, … , *N*} , where *n* = 0 denotes the fully unbound configuration (Fig. 2b).

**Figure 2.**
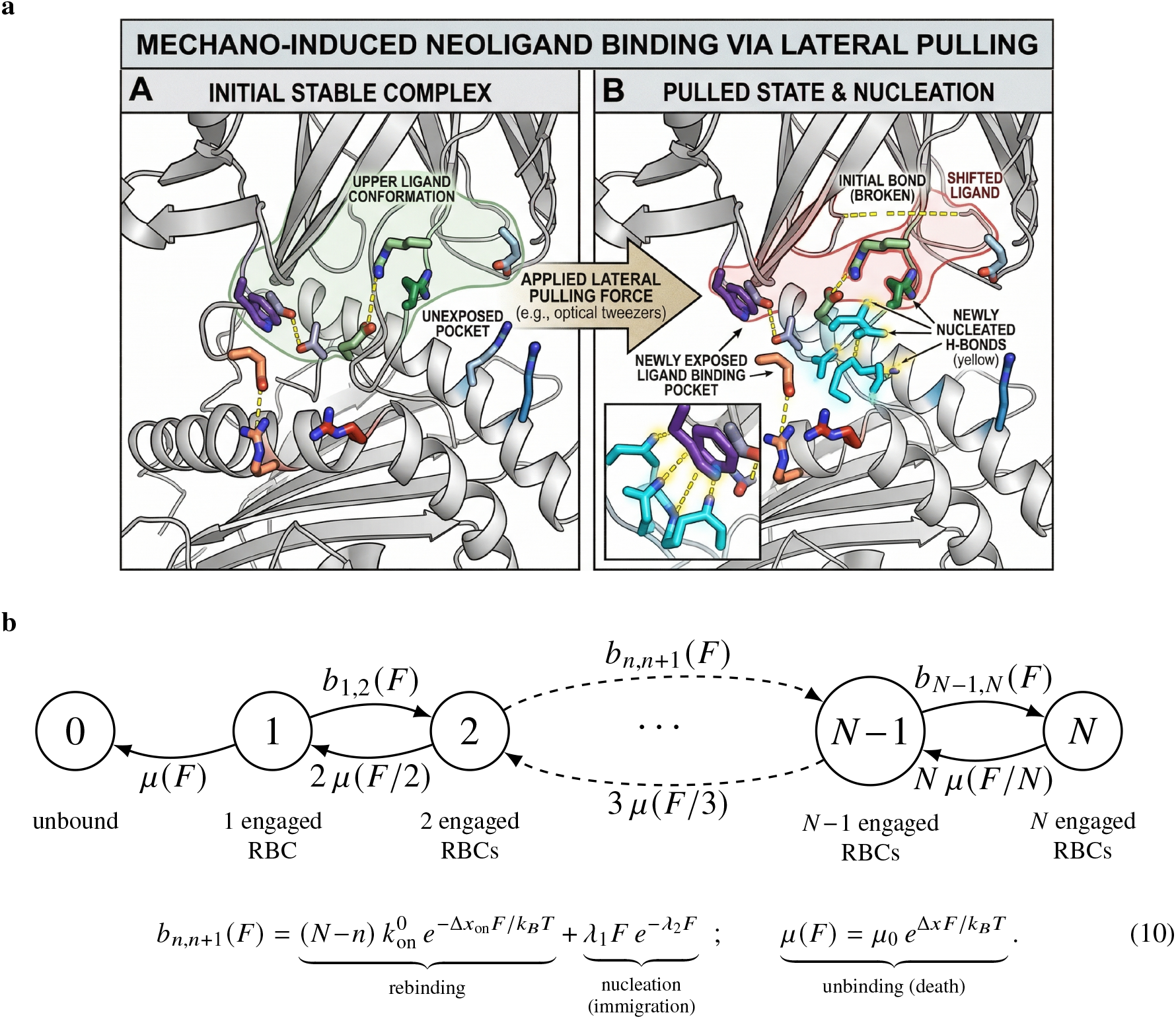
Molecular cartoon and birth–death description of the CFPR mechanism. **a)** Molecular schematic of mechano-induced nucleation. Lateral pulling (*e*.*g*., by optical tweezers) unzips the original upper ligand conformation and exposes an otherwise hidden ligand-binding pocket, whose newly exposed residues (orange, lysine; blue, arginine) nucleate new, stronger hydrogen bonds (yellow). Force therefore *recruits* additional residue binding contributions (RBCs), rather than only rupturing the existing ones —the molecular basis of the immigration term *λ* (*F*) (inspired by MD simulations in Ref. (29)). **b)** Coarse-grained birth–death ladder for a loaded TCR interphase. The state *n* counts the number of simultaneously engaged RBCs. RBCs are lost at the load-sharing death rate *d*_*n,n*−1_(*F*) = *n µ*(*F*/*n*), where each of *n* RBC shares the total force and senses *F*/*n*; the per-bond unbinding rate is given by Bell’s expression: 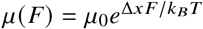 . RBCs are gained at the total birth rate *b*_*n,n*+1_(*F*) (Eq. (10)), which combines standard rebinding (prefactor 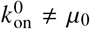, full force *F*) and force-induced immigration *λ*(*F*); the different prefactors reflect independent Kramers attempt frequencies for engagement and rupture. Rebinding is a forming event in which a single new RBC must overcome the full applied force *F*; load-sharing applies only to the *n* bonds already engaged. The effective distance Δ*x* can depend on pulling geometry, so changes in pulling angle modulate both engagement and loss of RBCs.

The Markov chain tracks the number of RBCs simultaneously engaged; the bond ruptures when all RBCs disengage. CFPR is powered by the irreversible mechanical work of force during RBC rupture and interphase reorganization. Pulling geometry can change molecular alignment and promote the engagement of additional RBCs, yielding a sharp crossover from slip-like to catch-like behavior at the *global* interphase lifetime. We focus on TCR-pMHC interactions, but the analysis applies to other receptor-ligand systems, including B-cell receptors in germinal centers (18).

Transitions between adjacent states follow a force-dependent total birth rate *b*_*n,n*+1_ (*F*) which includes a force-penalized rebinding rate from the *N* − *n* available sites, and a force-induced immigration rate, respectively,

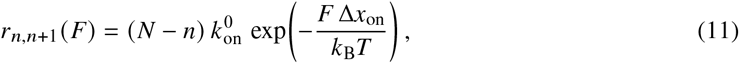

and

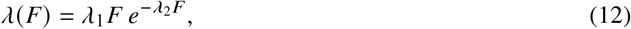

where 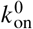 is the intrinsic zero-force association rate, Δ*x*_on_ is the rebinding distance parameter (Bell-like penalty), and *λ*_1_ and *λ*_2_ are parameters that quantify the force-induced amplification/deflation of the nucleation rate. The total birth rate entering simulations and likelihood evaluation is then

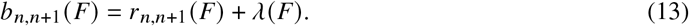

This total rate reduces to *λ*(*F*) when 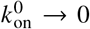 or at forces where immigration dominates rebinding but as shown below, the overall idea is preserved: microscopically, the catch-slip crossover is a consequence of the competition between a force-increasing unbinding of individual RBCs sharing the total load and a force-increasing recruitment of new RBCs through interphase reorganization. Similarly, we define a force-dependent, per-RBC death rate:

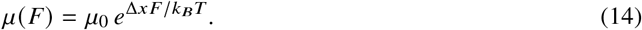

Unlike typical single-bond models, the experimental scenario imposes strain (e.g., micropipette displacement), and *F* is the resulting total force shared among all RBCs; hence, the load-sharing death rate has the form

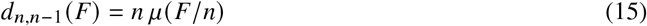

As shown below, catch-like behavior emerges in this model from *collective drift* of the multi-RBC state, not from catch behavior of any single RBC; each individual RBC follows slip-like (Bell/Kramers) dissociation kinetics.

Note that this is also a continuous-time Markov chain, so it predicts the multiexponential decay in the survival function, but it departs from other formulations in the fact that it can also use additional information from molecular dynamics and make falsifiable predictions about the presence or absence of the catch-bond effect, as we show in the following sections and in the Supplementary Material.

### 4.2 CFPR with constant rates (mean-field approximation)

We define the random variable *T*_*n*_ as the unbinding time if the ligand and receptor started with a bond with *n* RBCs. In section S3.2, we derive an exact expression for the mean value of this random variable, *τ*_*n*_ (*F*).

The equation for *τ*_*n*_ is cumbersome and not analytically tractable, so we evaluate it numerically below. However, to illustrate the implications of the model, we first analyze a simplified version obtained by introducing a mean-field approximation that replaces the *n*-dependent death rate *d*_*n,n*−1_(*F*) = *n µ*(*F*/*n*) with a constant per-state death rate *µ*(*F*) (Appendix S3). This approximation is not only analytically tractable but is valid when the number of engaged RBCs is small (*n ≪ N*), and the force is not too large, so that the load-sharing effect is weak. We note that while this approximation simplifies the analysis, it retains the essential physics of force-driven cooperative engagement and disengagement in the full CFPR model. Similarly, we replace the total birth rate *b*_*n,n*+1_ (*F*) with a state-independent birth rate *b* (*F*) that captures the average effect of force on RBC recruitment. Under these approximations, the mean bond lifetime satisfies the following simple linear relation:

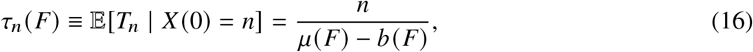

where (*X* (*t*)) _*t* ≥0_ represents the number of RBCs modeled by the CTMC in terms of the birth and death rates. As the bond is eventually broken, this approximation makes sense provided

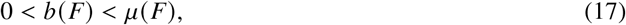

namely, that eventually the ligand and receptor will disengage.

Let us consider a TCR-pMHC interphase with *n* initially engaged RBCs (0 < *n ≪ N*). Equation (16) separates the *initial cooperative state n* from the *force dependence* in *µ* (*F*) − *b* (*F*) , so that CFPR produces non-trivial bond lifetime–force behavior when this drift is non-monotone in *F* —*i*.*e*., when force simultaneously increases the death rate and enables additional RBC immigration.

The maximum engagement time is attained when *dτ*_*n*_ (*F*)/*dF* = 0, which implies:

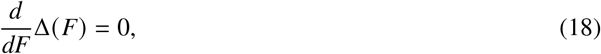

where Δ (*F*) ≡ *µ* (*F*) − *b* (*F*) is the net drift. This condition defines the catch-slip crossover and requires that, for some time, birth rates increase as the applied force increases. This is the case only if one postulates that rebinding is not proportional to 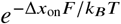 , and that force promotes additional RBC engagement through bond interphase reorganization. Several proposals, again based on energy landscape arguments, have been made to explain this effect (30, 31). Still, they are not based on a clear molecular mechanism (they replace the exponential dependence of the rebinding rate on *F* with a more complex function of *F*). In contrast, our CFPR mechanism provides a clear molecular explanation for this effect: force promotes additional RBC engagement by aligning the interphase and exposing new binding sites (Fig. 2a), where lateral pulling unzips the bound conformation and uncovers a hidden pocket whose residues nucleate new contacts.

In addition to Eq. (18), the catch-slip crossover also requires that the birth rate *b* (*F*) grows faster than the death rate *µ* (*F*) at low forces, but eventually the death rate dominates at high forces.To illustrate why our *nucleation* mechanism actually solves this problem, we introduce a simple parametrization of *µ* (*F*) and *b* (*F*) that captures two generic effects: force increases the death rate, while force and geometry promote interphase reorganization and additional RBC immigration. The loss rate grows exponentially with load (Bell/Kramers form),

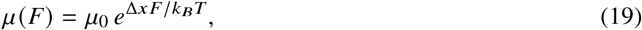

with 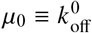 the zero-force off-rate and Δ*x* the transition-state distance for bond rupture. For the engagement rate, and using the mean-field approximation again, we use a minimal linear-damped form:

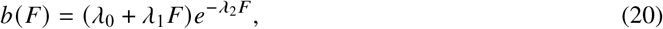

with *λ*_0_ ≥ 0 (s^−1^), *λ*_1_ ≥ 0 (s^−1^pN^−1^), and *λ*_2_ (*F*) ≥ 0 (pN^−1^).

This form provides a simple, effective description of rebinding (*λ*_0_; this is a conservative estimate of the rebinding rate to make the argument more clear-cut), nucleation (*λ*_1_), and high-force suppression (*λ*_2_). The linear term *λ*_1_*F* reflects the idea that force can promote additional RBC engagement by aligning the interphase and exposing new binding sites. The exponential suppression term 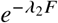 captures the structural limit on force-induced reorganization, preventing the engagement rate from growing unboundedly at high forces.

In our proof-of-concept simulations, we assume *λ*_0_ = 0 (Table S1), which represents a worst-case scenario for testing our mechanism (in Sec. S4.1 and the Supplementary Material, we show that the analysis is general across several plausible mathematical forms).

Let us assume, for the sake of simplicity, a linearized approximation of Eq. (20) (the engagement rate), such that *λ*_0_ = *λ*_2_ = 0 (no high-force suppression) and *b* (*F*) = *λ*_1_*F* (this is an overoptimistic scenario that makes the catch-slip crossover more pronounced, but we will show the most conservative case below). In this case, the binding time attains a maximum when:

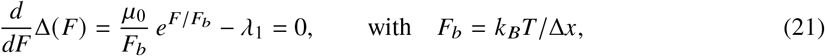

that is, at

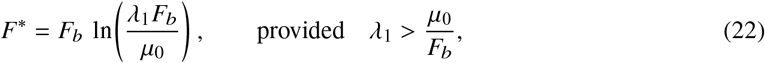

The condition *λ*_1_ > *µ*_0_ *F*_*b*_ holds for our parameters in Table S1 (0.08 s^−1^pN^−1^ > 0.1 8.28 s^−1^pN^−1^) yielding *F*^*\**^ *≃* 15.7 pN. For the full high-force–suppressed form (Eq. (20) with *λ*_2_ > 0), the condition *d*Δ (*F*)/ *dF* = 0 becomes

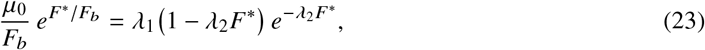

a transcendental equation without a closed-form solution. Numerical evaluation with our catch parameters (*λ*_2_ = 0.08 pN^−1^) gives *F*^*\**^ ≈6.1 pN, substantially closer to the simulation peak (≈5 pN) than the linearized estimate. The residual gap reflects the additional effect of load-sharing in the full classical-trajectory Monte Carlo (CTMC) simulation not captured by the mean-field *τ*_*m*_. In the linearized version of the engagement rate, the second derivative at 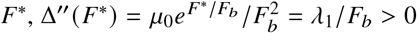, is positive; therefore, Δ (*F*) has a local minimum and *τ*_*m*_ (*F*) has a local maximum at *F*^*\**^. Note that the exact load-sharing model (Appendix S3) requires the strictly stronger *λ*_1_ > 2*µ*_0_ *F*_*b*_ for *τ* to be initially increasing at *F* = 0. Still, as the parameters need to be inferred from data, this is not a practical constraint for the proof-of-concept simulations, but it shows that nucleation is required and must be sufficiently strong to overcome the exponential loss of RBCs at low forces.

Equation (18) predicts a *sharp crossover*: at low forces (*F ≪ F*_*b*_), exponential *µ* (*F*) dominates and the interphase behaves slip-like; at intermediate forces, *λ* (*F* growth transiently reduces *µ* (*F*) −*λ* (*F*) , producing an emergent catch-like increase of *τ*_*m*_ (*F*) ; at larger forces, loss again dominates. This slip→catch→slip crossover is a collective effect of the CFPR ladder — a force- and geometry-driven bifurcation in effective interphase stability.

The extent of the catch regime across parameter space is summarized in Fig. 3, which maps the sign of *dτ*_*m*_ /*dF*|_*F*=0_ (catch when positive) computed from the exact absorbing CTMC. The mechanism is controlled by the competition between force-suppressed RBC loss and force-enhanced recruitment: catch requires slow death together with strong, persistent immigration (panels a, c), but is negligibly broadened by multivalency *N* (panel d). Finally, it is essentially independent of standard rebinding 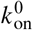 (panel b) — the same parameter that the inference below finds non-identifiable. Catch is thus a broad, robust regime rather than a marginal one. The complete set of fifteen two-parameter sweeps is given in the Supplementary Material (Sec. S7).

**Figure 3.**
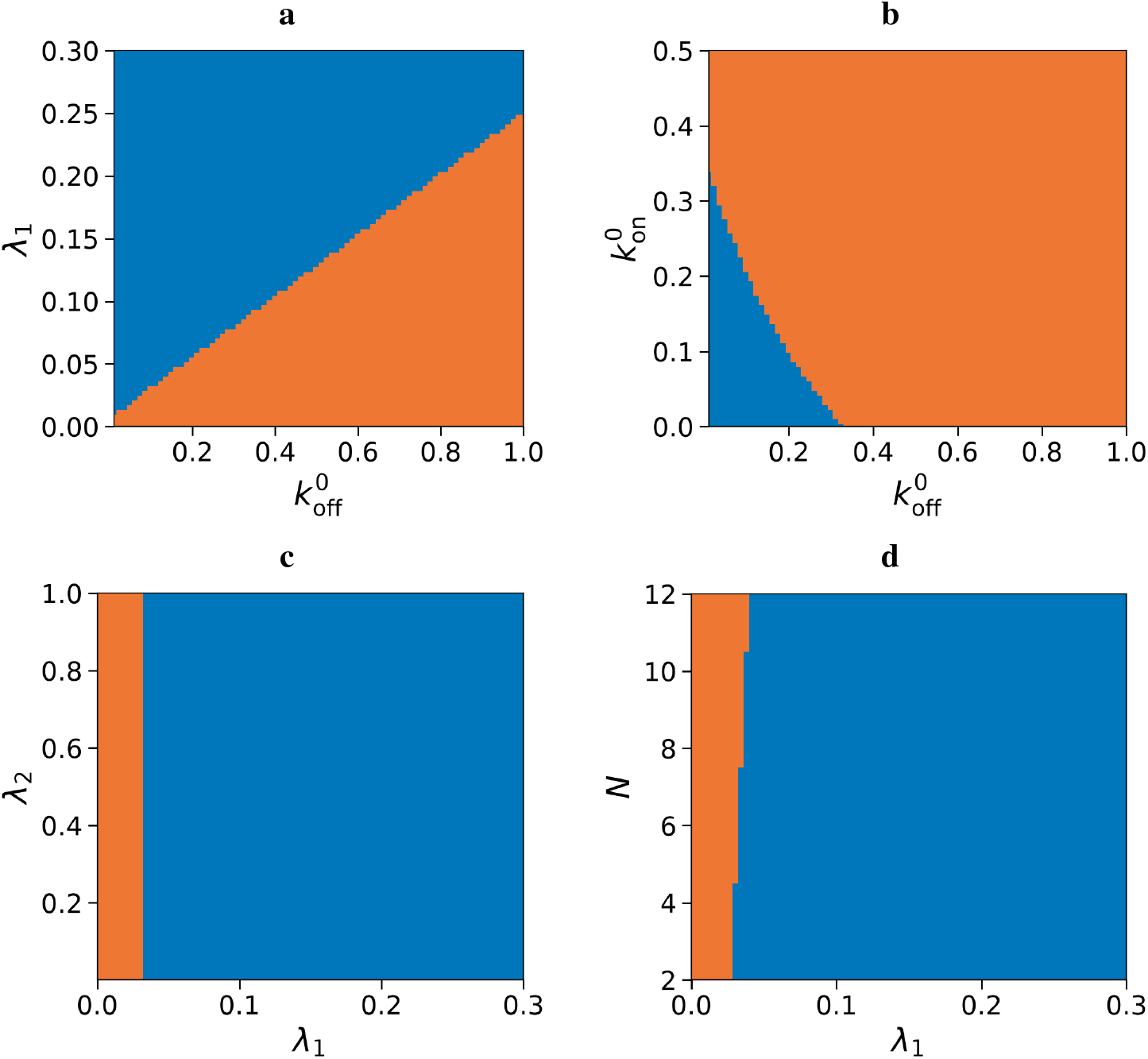
Catch/slip phase diagrams of the CFPR model. Each panel shows the sign of *dτ*_*m*_/ *dF*| _*F*=0_ from the exact absorbing-CTMC mean first-passage time over two swept parameters (others fixed at the catch-regime defaults, Table S1): 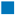 catch regime (> 0), 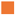 slip regime (≤ 0). **a)** death rate 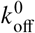 vs. immigration slope *λ*_1_: catch requires slow death and strong recruitment (approximately hyperbolic boundary). **b)** 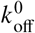 vs. rebinding 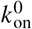 : the boundary is set by 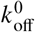 and is nearly independent of 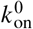 . **c)** immigration parameters *λ*_1_ vs. decay *λ*_2_: catch requires *λ*_1_ > 0 and slow high-force suppression. **d)** *λ*_1_ vs. valency *N*: more RBCs lower the immigration threshold for catch. See section S7 in the Appendix for the full set.

The CFPR model differs from the TP and TSTP models in two essential ways. First, it involves many internal states and emphasizes collective load sharing and recruitment rather than switching between a few conformations. Second, catch-like behavior arises from competition between two force-dependent processes — RBC loss and recruitment — whose balance depends on pulling geometry (Δ*x*) and reorganization (*λ* (*F*)). This provides a distinct route to catch bonding aligned with TCR interphase mechanics and highlights which effective force- and geometry-dependent functions would be most informative to estimate from molecular dynamics simulations.

Finally, we emphasize that CFPR is an out-of-equilibrium mechanism that also has a functional *discriminatory* effect. In particular, the mechanical work *W* =∫ *F dx* done by the external force powers both bond rupture and interphase reorganization. This is not simply a force-modified equilibrium: each RBC breakage dissipates energy through solvent friction, and interphase reorganization follows a non-equilibrium trajectory. Dissipation, on the other hand, can be quantified by the entropy production per cycle of unbinding-nucleation, which is positive for CFPR. Thus, entropy production (32) per forward cycle is

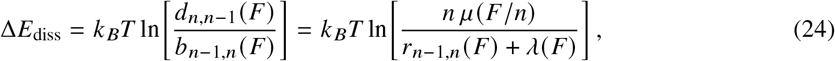

The return path requires mechanical work *W*_in_ ≥ Δ*E*_diss_. Since *d*_*n,n*−1_ > *b*_*n*−1,*n*_ when the drift toward disengagement is positive, Δ*E*_diss_ > 0 and net dissipation per cycle distinguishes CFPR from passive force-modified binding. Like molecular motors, CFPR is driven by mechanical work rather than ATP but shares the structure of out-of-equilibrium proofreading.

The CFPR mechanism also acts as a kinetic filter, unlike standard bonds in catch bond models. Weak binders dissociate before force-induced reorganization can recruit additional RBCs; stronger binders survive long enough to access the stable cooperative state. So, our choice for the term “proofreading” refers to this *functional outcome* — enhanced specificity through force-dependent kinetic filtering.

## 5 ANALYSIS OF THE CFPR MODEL

### 5.1 Force–lifetime curves and CFPR crossover

We simulated the CFPR birth–death process (Section 4) under slip and catch regimes. In the slip regime (*λ*_1_ = 0), the mean bond lifetime decreases monotonically with force (Bell-type). In contrast, in the catch regime (*λ*_1_ = 0.08 s^−1^pN^−1^, *λ*_2_ = 0.08 pN^−1^), the mean bond lifetime–force curve peaks at *F* ≈ 5 pN, with peak-to-baseline ratio ≈35-fold (at *F* = 5 pN the Kaplan–Meier *restricted mean survival time* (KM-RMST) is ≈ 44 s, while at *F* = 60 pN the KM-RMST is ≈ 1.25 s; Fig. 4a), confirming a collective emergent catch–slip bond behavior from individually slip-like RBCs. Representative trajectories (Fig. 5) show prolonged dwells at intermediate forces, reflecting competition between force-enhanced recruitment and accelerated loss.

**Figure 4.**
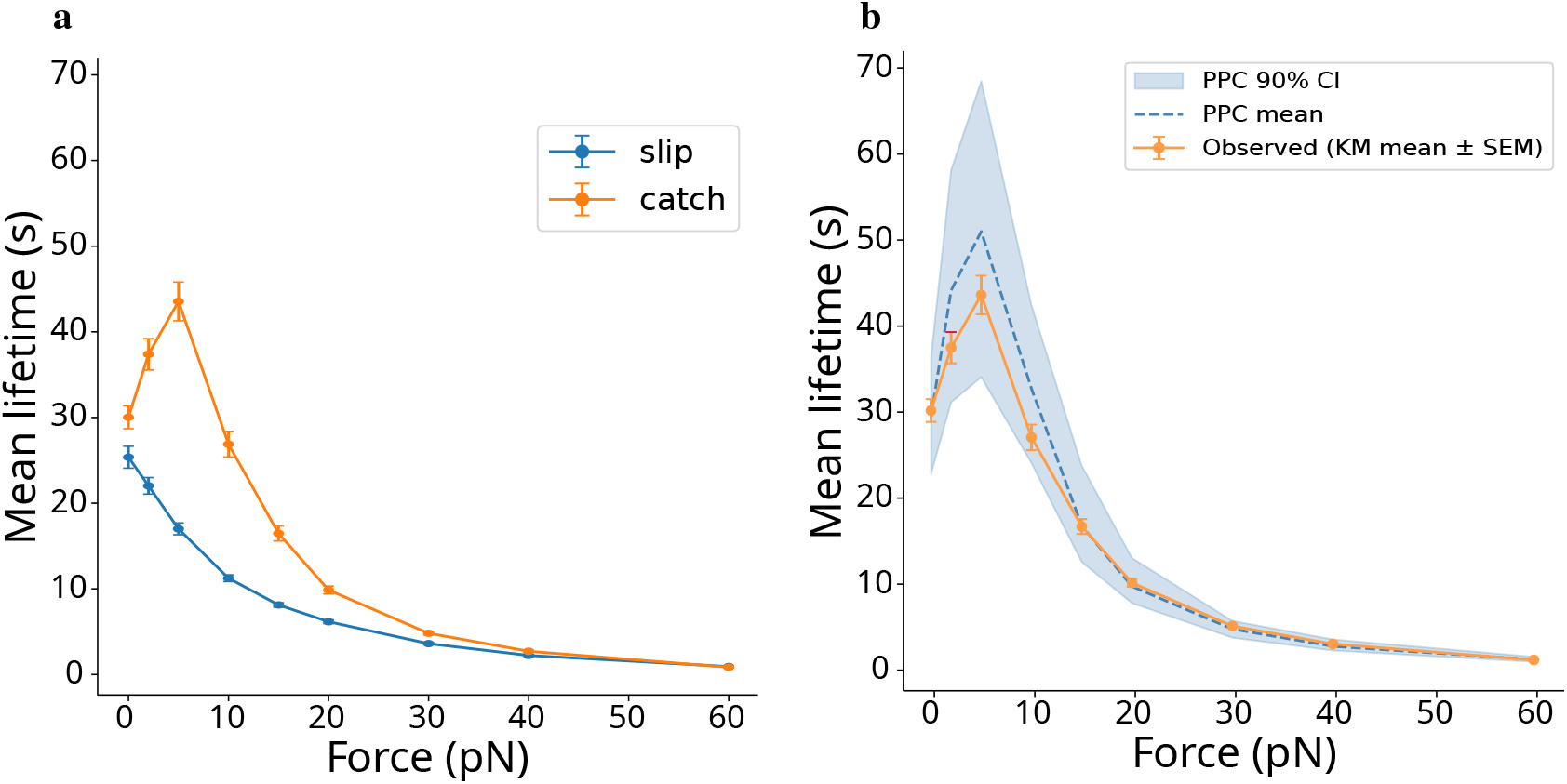
(a) Synthetic force–lifetime curves for slip and catch CFPR regimes. Mean bond lifetime (Kaplan–Meier RMST (KM-RMST), for *t*_max_ = 200 s) versus constant applied force. In the slip regime (*λ*_1_ = 0, *λ*_2_ = 0.1 pN^−1^) the KM-RMST decreases monotonically (blue circles), but in the catch regime (*λ*_1_ = 0.08 s^−1^ pN^−1^, *λ*_2_ = 0.08 pN^−1^) the KM-RMST follows a non-monotone catch-slip curve peaking at *F* ≈ 5 pN (KM-RMST ≈ 44 s) with peak-to-baseline ratio ≈ 35-fold (orange circles). Error bars: ±SEM. Parameters used are those indicated in Table S1. **(b) Posterior predictive check: force–lifetime curve**. Orange circles, observed KM mean lifetimes ±SEM (200 synthetic trajectories per force); these are the same values as the orange circles in panel (a). Blue band: 90% posterior predictive credible interval (100 posterior samples). Blue dashed line, mean of the posterior credible interval. All nine force values fall inside the 90% band (PPC coverage = 9/9), confirming the inferred model reproduces the observed catch-slip behavior.

**Figure 5.**
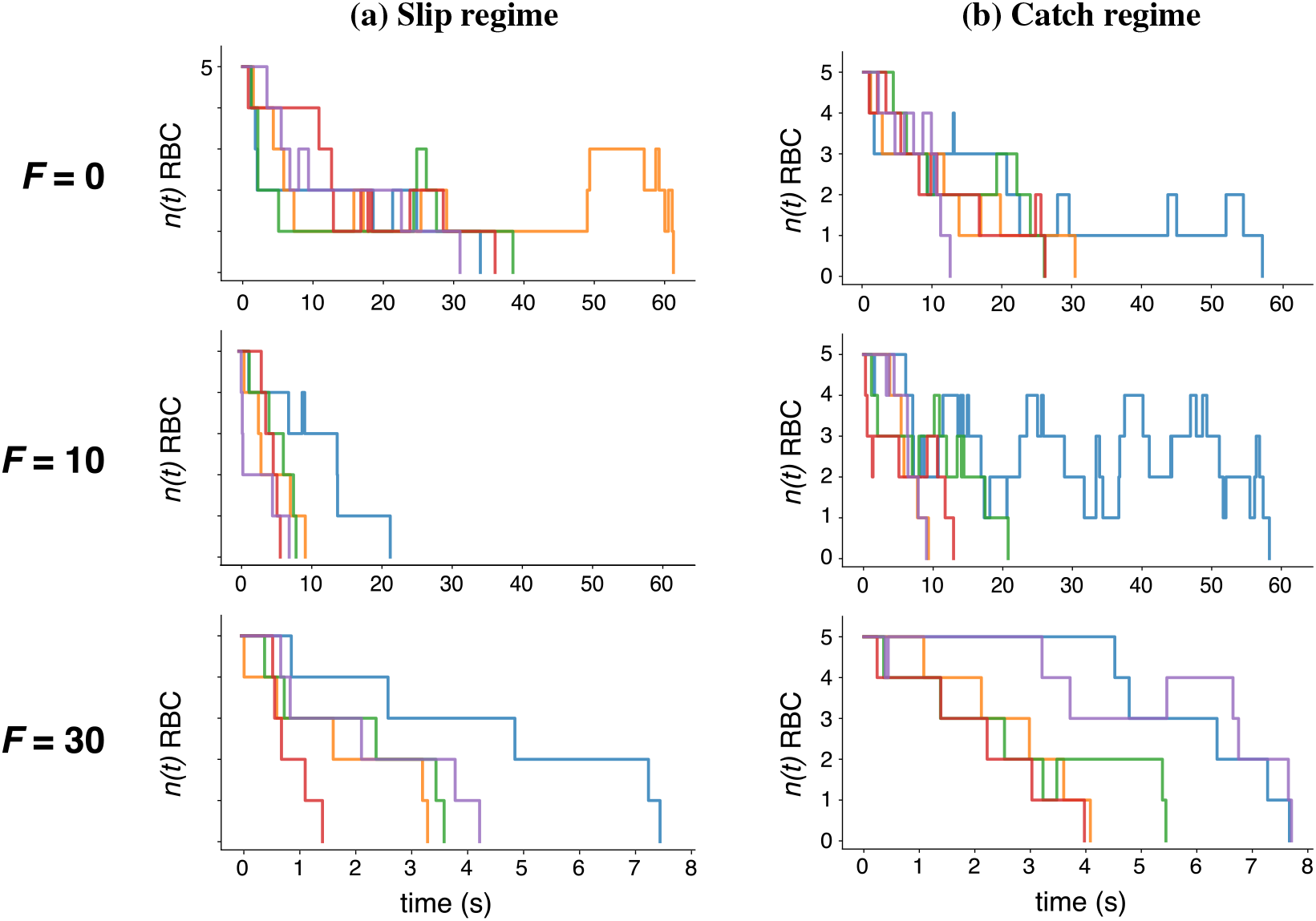
Representative CFPR trajectories under different parameter regimes. **(a)** Slip parameter regime. **(b)** Catch parameter regime. Number *n* (*t*) of RBCs at zero (*F* = 0 pN), intermediate (*F* = 10 pN, near the lifetime peak), and high (*F* = 30 pN) force. Trajectories terminate at absorption (*n* → 0). Longer dwell times at *F* = 10 pN and *F* = 30 pN in panel **(b)** (catch regime) reflect force-enhanced immigration *λ* (*F*). Also note the increased number of *birth* events (upward jumps) in **(b)** with respect to **(a)** at *F* = 10 pN and *F* = 30 pN.

### 5.2 Synthetic parameter recovery and identifiability

We performed an identifiability analysis with a synthetic experiment. We generated 200 CTMC trajectories at each of nine force values (*F ∈* {0, 2, 5, 10, 15, 20, 30, 40, 60} pN) under known ground-truth parameters and inferred all eight CFPR parameters by Hamiltonian Monte Carlo (Section S4.3). The dissociation parameters 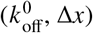 and immigration ones (*λ*_1_, *λ*_2_, *N*) were well identified: 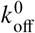 and Δ*x* recovered within 5% of true values (4% and 3%, respectively), and *N* exactly. The immigration parameters were recovered with larger mean bias: *λ*_1_ posterior mean 0.100 vs. true 0.080 (25% bias) and *λ*_2_ posterior mean 0.092 vs. true 0.080 (15% bias), though both true values fell within their 95% credible intervals (Table S3). The standard-rebinding parameters 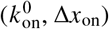 were poorly or not identified: Δ*x*_on_ posterior was nearly flat and the 95% CI for 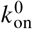 spanned [ 0.001, 0.019 ] s^−1^ (true 0.02 lies just above the upper bound). Force-induced immigration dominates the birth rate at 5–15 pN, masking standard rebinding’s force dependence. This is a concretely realizable failure mode even with fully observed trajectories. Identification of 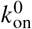 and Δ*x*_on_ requires independent zero-force measurements or protocol-dependent observables.

The posterior predictive check shows the inferred model is consistent with the observed catch-slip behavior across all nine force values (PPC 9/ 9, Fig. 4b; per-force predictive *p*-values 0.26–0.82, none near 0 or 1). The 90% credible band captures the presence of the peak in the mean lifetime, so the model is able both to capture the relevant parameters (Supplementary Material) while explaining the mechanism without relying just on fitting a survival function.

### 5.3 Bayesian diagnostics

Posterior marginals, trace plots, and pairwise correlations (Appendix S6) confirm that the dissociation and immigration arms are well identified (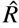 true values of 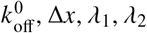 , and *N* all fall within their 95% credible intervals), while the two standard-rebinding parameters remain non-identifiable (Fig. S11). The full joint posterior (all pairwise marginals, Appendix S6.2) shows that the only strong correlations are the expected 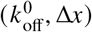 and (*λ*_1_, *λ*_2_) compensation ridges, showing that we have a broad set of plausible explanations. We do not need to rely on specific cherry-picked parameters to explain the catch-bond mechanism. A high-resolution posterior predictive check (*∼*10^4^ predictive lifetimes per force) reproduces not only the mean lifetime but the entire lifetime *distribution* at every force (Appendix S6.3), further supporting the main conclusion in the previous subsection.

This has an important biophysical implication. Force–lifetime trajectories — whether from single-molecule force spectroscopy or from loaded molecular-dynamics simulations of the contact zone — robustly constrain the quantities that set the catch behavior, namely, the intrinsic off-rate and transition-state distance of the load-bearing bonds 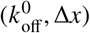 and the force-dependence of contact-zone recruitment (*λ*_1_, *λ*_2_). In contrast, they do *not* constrain the intrinsic association kinetics 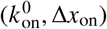 because, once force drives recruitment, the standard rebinding process contributes negligibly to the birth rate (Fig. 3b). Association parameters must therefore be supplied independently — from zero-force kinetic assays or structural measurements — rather than from force–bond lifetime curves.

## 6 DISCUSSION AND CONCLUSIONS

In this work, we show that the main mechanistic models of catch-slip bonds (Table 1) face similar non-identifiability problems and, because of that, cannot be distinguished from each other using only force-bond lifetime observables (16, 17). Consequently, discriminating among them requires complementary observables beyond curve fitting.

We quantified the validity domain of Kramers-type reductions via timescale separation, and find that the ratio *R* (*f*) = *τ*_relax_ /*τ*_escape_ remains well below 0.1 throughout the physiologically relevant 0–30 pN range for typical receptor–ligand parameters. However, quantitative predictions at very high forces (beyond *∼* 30 pN) warrant verification against a full Fokker-Planck or molecular dynamics treatment.

In view of that, and inspired by MD simulations in Ref. (29), we propose here a proofreading-type, cooperative mechanism driven by irreversible mechanical work (CFPR model) that provides a new approach to catch-like behavior. Sensitivity analysis confirms that the qualitative crossover of the catch-to-slip transition is robust with respect to alternative *λ* (*F*) functional forms, while highlighting the need for molecular-scale constraints.

Protocol-dependent force spectroscopy —comparing constant-force, force-ramp, and force-cycling measurements— can reveal hysteresis and loading-rate effects that rule out simple single-pathway descriptions, providing direct evidence of complex energy landscapes with multiple dissociation routes. Measuring effective off-rates with sub-piconewton force resolution (biomembrane force probes or optical tweezers (33, 34)) can detect deviations from Bell-like linearity in semi-log plots. For instance, the characteristic curvature predicted by the sliding-rebinding model, which arises from ratios of force-dependent dissociation Bell rates rather than simple exponentials, offers a concrete experimental discriminator between SR and TSTP or AD mechanisms.

Pulling geometry-dependent assays are particularly relevant to the CFPR mechanism proposed here. If synapse reorganization is geometry-sensitive, bond lifetimes should vary systematically with the direction of the applied force, yielding a signature that would distinguish network-level cooperative effects from intrinsic molecular catch bonding. Micropipette aspiration with controlled pulling angles can test this prediction directly (35). At the single-molecule level, combining force measurements with FRET or other structural reporters could detect conformational switching in real time, constraining the timescales of transitions that are currently hidden from bond-lifetime data alone. Fitting full survival curves *S* (*t, F*) to multi-exponential forms and tracking how weights *ω*_*i*_ and rates *k*_*i*_ evolve with force would further dissect the underlying pathway complexity. Encouragingly, the parameters in our model that *are* identifiable are precisely those that govern whether a given receptor–ligand interface will exhibit catch behavior, so trajectory-level data remain predictive for the property of interest.

Finally, molecular simulation of receptor–ligand complexes under load can provide critical constraints that are currently unavailable (15, 36). By estimating intrawell relaxation times and identifying the force threshold at which Kramers-type reductions break down, such simulations would bridge the gap between coarse-grained kinetic models and molecular-scale mechanics, grounding phenomenological parameters in physically measurable quantities. In particular, steered or constant-force simulations could directly test the mechano-induced nucleation scenario of Fig. 2a predicted by the CFPR model —that is, test whether lateral pulling exposes hidden binding pockets and nucleates new contacts— and so provide a solid basis to estimate the immigration rate *λ* (*F*) from first principles. Together, these approaches can transform the identifiability challenge from a limitation into an opportunity: rather than fitting curves, the field can design experiments that actively discriminate between mechanistically distinct conceptual schemes. Embedding validity-domain diagnostics and identifiability analyses into model fitting will help avoid overinterpretation of mechanistic conclusions from bond-lifetime curves.

More broadly, our unifying view supports a shift from static lymphocyte receptor-ligand affinity to a mechanosensitive description in which force dependence and cooperativity can be genuine targets of lymphocyte receptor selection.

## AUTHOR CONTRIBUTIONS

J.I. and M.C. implemented the computational pipeline and performed all numerical simulations and Bayesian inference. M.G.-S. contributed to the initial biological framing. J.F. and M.C. designed the research, formulated the CFPR mechanism, and wrote the manuscript. All authors reviewed and approved the final version.

## 7 ACKNOWLEDGMENTS

This work has been partially funded by Grant PID2022-140217NB-I00 and PID2025-169365NB-I00 funded by MCIN/AEI/ 10.13039/501100011033 (JI, MC), by Grant PRE2020-092274 funded by MCIN/AEI/

10.13039/501100011033 and by “ESF Investing in your future”, and by Xunta de Galiza under project GRC-ED431C 2020/02 (JF).

## DATA AND CODE AVAILABILITY

All simulation, inference, and visualization code is deposited at `[GitHub URL]`.

## SUPPLEMENTARY MATERIAL

An online supplement to this article can be found by visiting BJ Online at http://www.biophysj.org. The supplement includes: (1) Sensitivity analysis of CFPR with alternative *λ* (*F*) functional forms. (2) Full lifetime distribution analysis. (3) Analysis of effective rates for the sliding-rebinding model showing non-Bell-like behavior. (4) Detailed parameter tables for all simulations.

## Notes

### Competing Interest Statement

The authors have declared no competing interest.

