## Supplementary Material for "Emergent catch bonds from slip-like contacts: cooperative force-driven proofreading at receptor–ligand interfaces"

#### S1 VALIDITY OF PHENOMENOLOGICAL MODELS

Diffusion in an energy landscape involves two timescales: fast relaxation within a well ($\tau_f$) and slow escape over a barrier ($\tau_s$). Phenomenological models capture only the slow dynamics via master equations that describe transitions between metastable states. Their validity requires $\tau_s \gg \tau_f$, so the distribution within each well adjusts rapidly to slow changes in occupancy.

This separation is central to deriving master equations from the Fokker–Planck or Smoluchowski description (37, 38). Below, we review Kramers' steady-state current method for computing escape rates, which forms the basis of the Arrhenius–Kramers expression used in most of the coarse-grained models analyzed here.

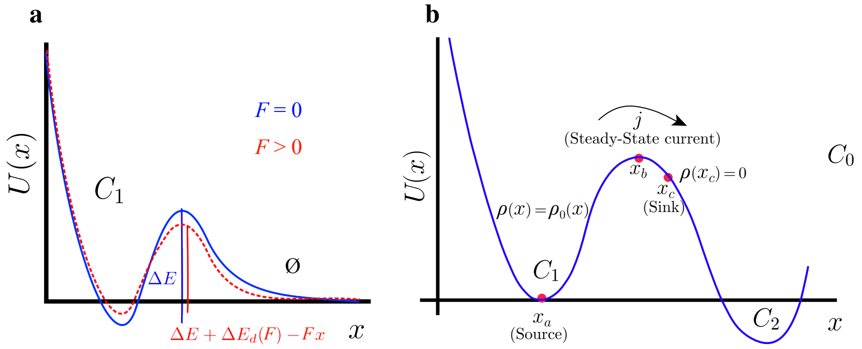


**Figure S1. Free-energy landscapes and Kramers' escape. a)** Beyond linear Bell tilting ($-Fx$), force-induced deformation energy $\Delta E_d(F)$ raises the effective barrier at intermediate loads before the slip pathway dominates at high force. **b)** Kramers' steady-state current. A source at $x_a$ feeds particles into the bound well; a sink at $x_c > x_b$ absorbs particles crossing the barrier at $x_b$. The steady-state current $j$ (in general, a function of the probability density $\rho(x,t)$, from Eq. (S2)) is proportional to the unbinding (escape) rate $k_{\text{off}}$ (Eq. S1).

##### Kramers' rate escape formula

Consider an energy landscape with a source at $x_a$ and a sink at $x_c > x_b$ (barrier top) like that in Fig. S1b. Because $\tau_s \gg \tau_f$, particles equilibrate within the well before escaping, so a quasi-stationary current $j$ flows from $x_a$ toward $x_c$. This current is related to the unbinding (escape) rate $k_{\text{off}}$ and the equilibrium well population $n_1$ by

$$j = k_{\text{off}} \times n_1. \tag{S1}$$

We obtain $j$ and $n_1$ from the Smoluchowski equation, the high-friction (overdamped) limit of the Fokker–Planck equation appropriate for molecular processes in aqueous solution, written as a continuity equation for the probability density $\rho(x,t)$ with probability current $j(x,t)$:

$$\frac{\partial \rho(x,t)}{\partial t} = -\frac{\partial}{\partial x} j(x,t), \qquad j(x,t) = -\frac{1}{\eta}\left[U'(x)\,\rho(x,t) + k_b T\,\frac{\partial \rho(x,t)}{\partial x}\right], \tag{S2}$$

with $\eta$ the drag coefficient. In steady state $\partial\rho/\partial t = 0$, so $\partial j/\partial x = 0$ and the current $j$ is spatially constant. The flux relation in Eq. (S2) then becomes a first-order linear ODE for $\rho(x)$,

$$U'(x)\,\rho(x) + k_b T\,\rho'(x) = -\eta j, \tag{S3}$$

which we integrate using the integrating factor $e^{U(x)/k_bT}$:

$$\frac{d}{dx}\left[e^{U(x)/k_bT}\,\rho(x)\right] = -\frac{\eta j}{k_b T}\,e^{U(x)/k_bT}. \tag{S4}$$

Integrating from $x_a$ to the absorbing sink at $x_c$ and imposing $\rho(x_c) = 0$ gives

$$e^{U(x_a)/k_bT}\,\rho(x_a) = \frac{\eta j}{k_bT}\int_{x_a}^{x_c} e^{U(x)/k_bT}\,dx. \tag{S5}$$

The well population is set by the quasi-equilibrium distribution $\rho_0(x) = \mathcal{N}\,e^{-U(x)/k_bT}$ near the minimum: normalizing over the well, $\int_{-\infty}^{x_b}\rho_0(x)\,dx = n_1$, fixes the left-hand side as $e^{U(x_a)/k_bT}\rho(x_a) = n_1\big/\int_{-\infty}^{x_b} e^{-U(x)/k_bT}\,dx$. Combining this with Eq. (S5) and using $k_{\text{off}} = j/n_1$ yields Kramers' unbinding rate:

$$k_{\text{off}} = \frac{k_bT}{\eta}\left[\int_{x_a}^{x_c}\exp(U/k_bT)\,dx\right]^{-1}\cdot\left[\int_{-\infty}^{x_b}\exp(-U/k_bT)\,dx\right]^{-1}. \tag{S6}$$

For $\Delta E \gg k_bT$, both integrals are dominated by narrow neighborhoods: the first by the barrier top $x_b$, the second by the well minimum $x_a$. Expanding the potential quadratically, $U(x) \approx U(x_a) + \frac{1}{2}U''(x_a)(x-x_a)^2$ near the well and $U(x) \approx U(x_b) - \frac{1}{2}|U''(x_b)|(x-x_b)^2$ near the barrier (where the curvature is negative), the resulting Gaussian integrals give

$$\int_{-\infty}^{x_b} e^{-U/k_bT}\,dx \approx e^{-U(x_a)/k_bT}\sqrt{\frac{2\pi k_bT}{U''(x_a)}},\qquad \int_{x_a}^{x_c} e^{U/k_bT}\,dx \approx e^{U(x_b)/k_bT}\sqrt{\frac{2\pi k_bT}{|U''(x_b)|}}. \tag{S7}$$

Substituting into Eq. (S6) and writing $\Delta E = U(x_b) - U(x_a)$ for the barrier height, the prefactors of $k_bT/\eta$ combine into the following Arrhenius–Kramers form of the zero-force escape rate:

$$k_{\text{off}} \approx \frac{\omega_A\omega_B}{2\pi\eta}\exp(-\Delta E/k_bT), \tag{S8}$$

with $\omega_A = [U''(x_a)]^{1/2}$ and $\omega_B = [|U''(x_b)|]^{1/2}$. Since the above equation is essentially the same as Eq. 2, the prefactor $\mu$ in this last equation is, therefore, $\mu = \omega_A\omega_B/(2\pi\eta)$, that is, the attempt frequency at which the bound complex rattles against the barrier, while $e^{-\Delta E/k_bT}$ is the probability of a thermal fluctuation large enough to surmount it. For a typical receptor–ligand barrier $\Delta E \approx 15\,k_bT$, $e^{-15} \approx 3\times10^{-7}$, so escape is rare and the timescale separation $\tau_s \gg \tau_f$ is self-consistent.

The effect of force enters through the landscape itself. An applied tensile force $F$ along $x$ adds a mechanical potential $-Fx$, so $U(x,F) = U_0(x) - Fx$. To leading order, this leaves the extrema positions fixed but lowers the barrier height to $\Delta E(F) = \Delta E_0 - F\,\Delta x$, where $\Delta x = x_b - x_a$ is the well-to-barrier distance. Inserting this into Eq. (S8) factorizes the exponential and recovers the Bell form (Eq. 4), $k(F) = k^0\,e^{F/F_b}$ with Bell force scale $F_b = k_bT/\Delta x$, so the exponential increase of the off-rate with force is the slip-bond baseline against which catch behavior is measured.

As mentioned in the main text, Sec. 3, Bell-type descriptions thus assume force affects only barrier heights, not well/barrier positions or curvatures. Their validity requires (i) $\Delta E \gg k_bT$ (ensuring $\tau_s \gg \tau_f$) and (ii) barriers sharp enough that extrema positions depend weakly on force (9). Condition (i) can be made quantitative: intra-well relaxation $\tau_f \sim 1/\omega$ (with $\omega \approx 10^{12}\,\text{s}^{-1}$ for protein–ligand systems) must remain much faster than escape $\tau_s = 1/k(F)$, *i.e.*,

$$R(F) = \frac{\tau_f}{\tau_s} = \frac{k(F)}{\omega} = \frac{k^0}{\omega}\,e^{F/F_b} \ll 1. \tag{S9}$$

For $\Delta E_0 = 15\,k_bT$, $k^0/\omega = e^{-15} \approx 3\times10^{-7}$; even at $F = 30\,\text{pN}$ ($F/F_b \approx 3.6$ for $\Delta x = 0.5\,\text{nm}$) one finds $R \approx 10^{-5} \ll 1$. As shown in the main text (Section 3), the approximation holds throughout the physiological range, and condition (i) fails only when $\Delta E(F) < 5k_BT$ ($F > F_{\text{breakdown}} \approx 30\,\text{pN}$ for typical TCR parameters), necessitating full Fokker-Planck or molecular dynamics approaches.

#### S2 EXTENDED CATCH–SLIP BOND MODELS AND REDUCTIONS

Catch-bond behavior has been documented in selectins, integrins, bacterial adhesins, and T cell receptors, and several mechanisms have been proposed (5, 6, 39). These mechanisms map onto a small number of conceptual models (Table 1 in the main text). We review here these models as originally formulated and show that many are either indistinguishable at the level of typical observables or mathematically equivalent, posing a significant challenge for mechanism identification.

#### S2.1 Two-pathway model (TP)

The two-pathway model (9, 40) assumes a one-dimensional potential $U(x)$ along the reaction coordinate $x$. The landscape has a minimum at $x = 0$ (bound state) and two barriers at $x_c < 0$ (catch) and $x_s > 0$ (slip) (Fig. S2). An applied force tilts the landscape (Eq. 3), raising one barrier while lowering the other.

As shown in Appendix S1, this model reduces to a Markov chain via Kramers' theory. The bound state $C_1$ dissociates via two pathways:

$$\emptyset \xleftarrow{\;k_c(F)\;} C_1 \xrightarrow{\;k_s(F)\;} \emptyset \tag{S10}$$

where $\emptyset$ denotes the unbound state. In this case,

$$\frac{1}{\tau(F)} = k(F) = k_c^0 \exp\left(\frac{x_c F}{k_B T}\right) + k_s^0 \exp\left(\frac{x_s F}{k_B T}\right), \tag{S11}$$

with $k_c^0$ and $k_s^0$ the zero-force rates and $x_c$, $x_s$ the associated Bell lengths for the catch and slip barriers, respectively.

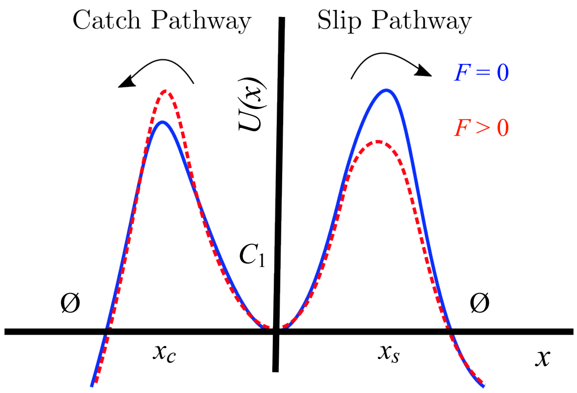


**Figure S2. Two-pathway model landscape.** The two-pathway model assumes a single bound minimum connected to the unbound state via two barriers, located at $x_c < 0$ (catch) and $x_s > 0$ (slip). Under a right-directed force, the catch barrier rises, and the slip barrier lowers, progressively favoring dissociation through the slip pathway.

This model captures catch-slip behavior when the catch barrier is lower than the slip barrier at zero force. As force increases, the catch barrier rises, and the slip barrier lowers. The mean lifetime increases until the two barriers become equal; beyond that, dissociation proceeds via the slip pathway. The scenario is mathematically straightforward but requires finely tuned barrier positions with no obvious structural interpretation (5, 6). Moreover, as shown in section S2.6, this model cannot reproduce the multi-timescale survival structure of Eq. (8).

#### S2.2 Two-state, two-pathway model (TSTP)

Many ligand-receptor systems can adopt multiple conformational states. In the main text, we have discussed the better-studied case of TCR, but this applies to other systems as well. From an immunological perspective, even Abs and BCRs can adopt multiple conformational states (25, 26, 41–43). The TSTP model (10, 11) extends the two-pathway framework with two bound states $C_1, C_2$ that interconvert with rates $k_{12}$ and $k_{21}$ and dissociate to the unbound state, respectively, with dissociation rates $k_{10}, k_{20}$ (Fig. S3), and leads to the coupled master equations for the state probabilities $P_1(t, F)$ and $P_2(t, F)$:

$$\begin{cases} \dfrac{dP_1(t,F)}{dt} = -\left[k_{12}(F) + k_{10}(F)\right] P_1(t,F) + k_{21}(F) P_2(t,F), \\ \dfrac{dP_2(t,F)}{dt} = k_{12}(F) P_1(t,F) - \left[k_{21}(F) + k_{20}(F)\right] P_2(t,F), \end{cases} \tag{S12}$$

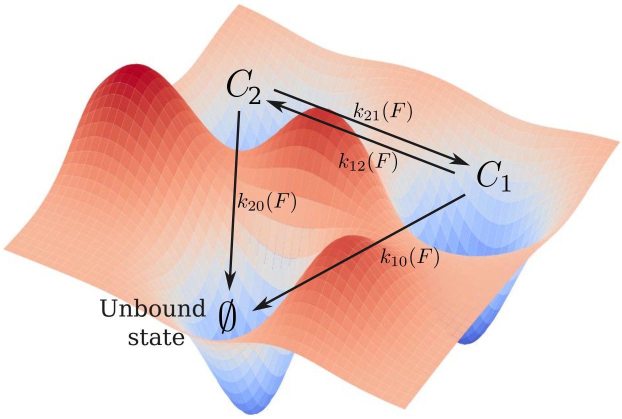


**Figure S3. High-dimensional free-energy landscape.** A receptor–ligand interaction is governed by an underlying high-dimensional free-energy landscape that can be projected onto a one-dimensional reaction coordinate $x$.

where each rate follows the Bell equation (4):

$$\begin{aligned} k_{10}(F) &= k_{10}^0 \exp\left(\frac{x_{10}F}{k_BT}\right), \quad k_{20}(F) = k_{20}^0 \exp\left(\frac{x_{20}F}{k_BT}\right), \\ k_{12}(F) &= k_{12}^0 \exp\left(\frac{x_{12}F}{k_BT}\right), \quad k_{21}(F) = k_{21}^0 \exp\left(\frac{x_{21}F}{k_BT}\right). \end{aligned} \tag{S13}$$

The system (S12) can be easily solved for fixed $F$, yielding:

$$\begin{cases} P_1(t,F) = \sigma_1 e^{-\lambda_1 t} + \sigma_2 e^{-\lambda_2 t}, \\ P_2(t,F) = \rho_1 e^{-\lambda_1 t} + \rho_2 e^{-\lambda_2 t}, \end{cases} \tag{S14}$$

where:

$$\lambda_1 = \frac{1}{2}\Big(k_{10} + k_{12} + k_{20} + k_{21} + \sqrt{(k_{10} + k_{12} + k_{20} + k_{21})^2 - 4(k_{10}k_{20} + k_{12}k_{20} + k_{10}k_{21})}\Big),$$

$$\lambda_2 = \frac{1}{2}\Big(k_{10} + k_{12} + k_{20} + k_{21} - \sqrt{(k_{10} + k_{12} + k_{20} + k_{21})^2 - 4(k_{10}k_{20} + k_{12}k_{20} + k_{10}k_{21})}\Big),$$

$$\sigma_1 = \frac{P_1(0)\Big(k_{10} + k_{12} - k_{20} - k_{21} + \sqrt{(k_{10} + k_{12} - k_{20})^2 + 2(-k_{10} + k_{12} + k_{20})k_{21} + k_{21}^2}\Big) - 2k_{21}P_2(0)}{2\sqrt{(k_{10} + k_{12} + k_{20} + k_{21})^2 - 4\big(k_{12}k_{20} + k_{10}(k_{20} + k_{21})\big)}},$$

$$\sigma_2 = \frac{P_1(0)\big(-k_{10} - k_{12} + k_{20} + k_{21}\big) + \sqrt{(k_{10} + k_{12} - k_{20})^2 + 2(-k_{10} + k_{12} + k_{20})k_{21} + k_{21}^2} + 2k_{21}P_2(0)}{2\sqrt{(k_{10} + k_{12} + k_{20} + k_{21})^2 - 4\big(k_{12}k_{20} + k_{10}(k_{20} + k_{21})\big)}},$$

$$\rho_1 = \frac{P_2(0)\Big(-k_{10} - k_{12} + k_{20} + k_{21} + \sqrt{(k_{10} + k_{12} - k_{20})^2 + 2(-k_{10} + k_{12} + k_{20})k_{21} + k_{21}^2}\Big) - 2k_{12}P_1(0)}{2\sqrt{(k_{10} + k_{12} + k_{20} + k_{21})^2 - 4\big(k_{12}k_{20} + k_{10}(k_{20} + k_{21})\big)}},$$

$$\rho_2 = \frac{P_2(0)\Big(k_{10} + k_{12} - k_{20} - k_{21} + \sqrt{(k_{10} + k_{12} - k_{20})^2 + 2(-k_{10} + k_{12} + k_{20})k_{21} + k_{21}^2}\Big) + 2k_{12}P_1(0)}{2\sqrt{(k_{10} + k_{12} + k_{20} + k_{21})^2 - 4\big(k_{12}k_{20} + k_{10}(k_{20} + k_{21})\big)}},$$

with $P_2(0) = 1 - P_1(0)$. We notice that, in those expressions, we have omitted, in all terms, the dependency on $F$ for better clarity.

Adding both equations in system (S14), we obtain the equation for the bond survival probability, $S(t,F) = P_1(t,F) + P_2(t,F)$:

$$S(t,F) = \omega_1 e^{-\lambda_1 t} + \omega_2 e^{-\lambda_2 t}, \tag{S15}$$

where $\omega_1 = \sigma_1 + \rho_1$ and $\omega_2 = \sigma_2 + \rho_2$; finally, it is straightforward to obtain $\omega_1 + \omega_2 = 1$.

The TSTP model reproduces catch-bond behavior when the two bound states differ in stability and force sensitivity. For instance, $C_1$ may be more populated at zero force but have a faster off-rate than $C_2$. Force shifts the equilibrium toward $C_2$ and/or differentially modulates the off-rates, so the effective lifetime increases with force over a range. This provides a kinetic representation of force-modulated conformational selection (5, 6).

Because two bound states with distinct off-rates contribute to dissociation, the survival probability $S(t, F)$ is generically *biphasic* rather than a single exponential (see a qualitative illustration of this biphasic decay in Fig. S4): a fast initial drop reflects depletion of the short-lived state $C_1$, followed by a shallow tail set by the long-lived state $C_2$, according to equation (S15), where $\lambda_{1,2}$ represent the fast and slow scales, respectively.

This is the simplest realization of the multi-exponential form of Eq. (8) (main text), with two modes. Increasing force shifts statistical weight toward the slow component and/or lowers its rate, lengthening the tail and prolonging the mean lifetime over a force range — the kinetic signature of catch behavior.

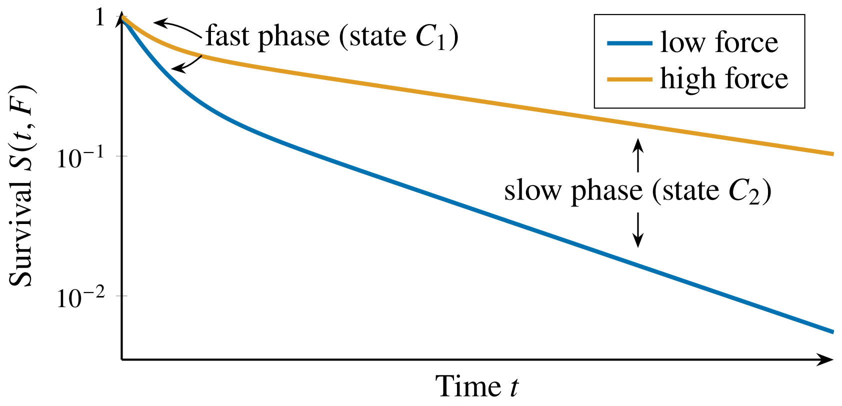


**Figure S4. Illustrative representation of a biphasic bond survival in the two-state, two-pathway model.** Schematic survival probability $S(t, F)$ on a semi-logarithmic scale for two applied forces. Two bound states with different off-rates produce a fast initial decay (depletion of the short-lived state $C_1$) followed by a shallow tail (long-lived state $C_2$); the change of slope is the hallmark of the two-mode form of Eq. (8). Increasing force shifts weight toward the slow component and reduces its rate, raising the tail and prolonging the mean lifetime — a catch-like response.

#### S2.3 Allosteric deformation model (AD)

Following Monod *et al.* (27), the AD model (13, 14) assumes two receptor conformations: $R_1$ (tense, low affinity) and $R_2$ (relaxed, high affinity). Force biases the receptor toward the relaxed state (5, 6). The key difference from the TSTP model is that conversion and dissociation are described by separate potentials (Fig. S5a,b). The model uses the bond survival probability $S(t, F)$ plus the probabilities $Q_1(t, F)$ and $Q_2(t, F)$ for the bound complex in states $C_1 = R_1 L$ and $C_2 = R_2 L$, respectively, such that $Q_1(t, F) + Q_2(t, F) = 1$ at all times and:

$$\begin{cases} \dfrac{dQ_1(t,F)}{dt} = -k_{12}(F)Q_1(t,F) + k_{21}(F)Q_2(t,F), \\[2ex] \dfrac{dQ_2(t,F)}{dt} = k_{12}(F)Q_1(t,F) - k_{21}(F)Q_2(t,F), \end{cases} \tag{S16}$$

The conversion rates are force-dependent and follow Bell-type expressions,

$$\begin{aligned} k_{12}(F) &= k_{12}^0 \exp\left(\frac{x_{12}F}{k_B T}\right), \\ k_{21}(F) &= k_{21}^0 \exp\left(\frac{x_{21}F}{k_B T}\right), \end{aligned} \tag{S17}$$

with $k_{12}^0$ and $k_{21}^0$ the zero-force rates and $x_{12}, x_{21}$ the associated Bell lengths.

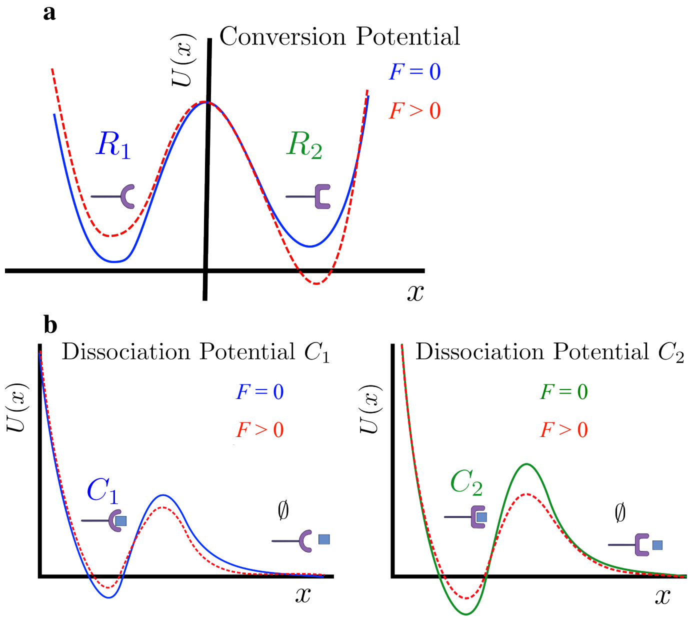


**Figure S5. Allosteric model landscapes. a)** Conversion potential. The receptor fluctuates between an inactive state $R_1$ and an active state $R_2$; force shifts the balance toward $R_2$. **b)** Dissociation potentials. Each state has its own dissociation potential, with a deeper well for the inactive state. Force reduces barrier heights and, by favoring $R_2$, can produce catch behavior.

Dissociation from each state is described by separate potentials $U_1(x)$ and $U_2(x)$ and corresponding off-rates,

$$\begin{aligned} k_{10}(F) &= k_{10}^0 \exp\left(\frac{x_{10}F}{k_B T}\right), \\ k_{20}(F) &= k_{20}^0 \exp\left(\frac{x_{20}F}{k_B T}\right), \end{aligned} \tag{S18}$$

where $k_{10}^0$ and $k_{20}^0$ are the zero-force dissociation rates and $x_{10}$, $x_{20}$ are the barrier widths for each potential. Because the receptor continuously fluctuates between $R_1$ and $R_2$, the bound complex experiences a time-dependent dissociation rate,

$$k(t, F) = k_{10}(F)Q_1(t, F) + k_{20}(F)Q_2(t, F), \tag{S19}$$

and the survival probability satisfies

$$\frac{dS(t, F)}{dt} = -k(t, F)\, S(t, F). \tag{S20}$$

In this framework, catch behavior arises when force both stabilizes the high-affinity state and differentially modulates the dissociation barriers, effectively prolonging the lifetime over a range of forces before slip-like behavior sets in.

However, while the underlying biophysical interpretation is different, note that Eqs. (S16) and Eqs. (S12) are mathematically identical, so they share the same solution for the survival function (see below subsection S2.6).

#### S2.4 Bond deformation model (BD)

The BD model (12) uses a single one-dimensional potential with one barrier (Fig. S1a). Force induces deformations of the binding pocket that strengthen or weaken non-covalent interactions. The dissociation rate includes an additional deformation energy $\Delta E_d(F)$:

$$k(F) = k_0 \exp\left[-\frac{\Delta E_d(F) - xF}{k_B T}\right], \tag{S21}$$

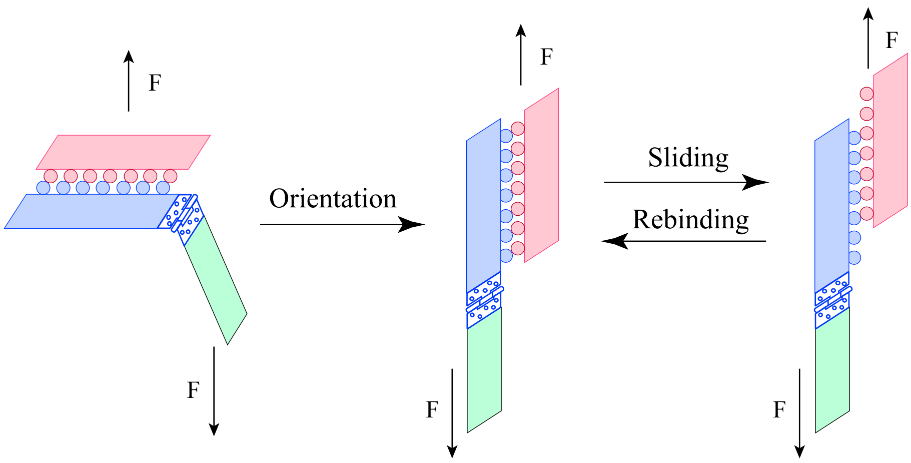


**Figure S6. Sliding rebinding.** Schematic representation of a receptor with a hinge domain that can slide relative to its ligand under force, breaking and reforming multiple non-covalent contacts before full dissociation.

where $-xF$ is the Bell tilting term. Pereverzev and Prezhdo (12) proposed:

$$\Delta E_d(F) = \begin{cases} \alpha\left[1 - \exp(-F/F_0)\right], & F < F_0, \\ \alpha, & F \geq F_0, \end{cases} \tag{S22}$$

with $\alpha$ the limiting deformation energy at threshold $F_0$. Catch behavior emerges when $\Delta E_d(F)$ dominates the Bell term over an intermediate force range, raising the effective barrier before slip-like behavior takes over.

#### S2.5 Sliding rebinding model (SR)

The models above are largely agnostic about the detailed structure of the receptor–ligand interface, so the reaction coordinates may lack a direct physical interpretation. The SR model (15) addresses this by proposing an explicitly structural mechanism for receptors with hinge-like domains: under force, the receptor slides relative to the ligand, transiently breaking contacts while forming new ones, enabling catch behavior through repeated rebinding.

The interface is represented by $N$ pseudoatom pairs that form identical non-covalent bonds, each with on-rate $k_{+1}$ and Bell-type off-rate $k_{-1}(F)$. When all $N$ bonds are broken, the complex can fully dissociate or slide to a new configuration where up to $N-1$ bonds can reform, with a force-dependent probability reflecting receptor orientation (Fig. S6).

For illustration, consider the case $N = 2$. The Markov-chain representation introduced in Ref. (15) can be reformulated in terms of four states, labeled $(ij)$, where $i$ denotes the maximum number of potential pseudoatom pairs and $j$ the number of bound pairs. The resulting network is shown in Fig. S7A, where transitions correspond to bond formation, bond breakage, and sliding:

The master equations for the probabilities $p_{ij}(t)$ of these four states are

$$\begin{cases} \dfrac{dp_{22}}{dt} = 2k_{+1}p_{21} + k_{+2}p_{11} - k_{-2}p_{22}, \\ \dfrac{dp_{21}}{dt} = k_{-2}p_{22} - 2\left(k_{+1} + p_n k_{-1} + (1-p_n)k_{-1}\right)p_{21}, \\ \dfrac{dp_{11}}{dt} = 2p_{\text{n}}k_{-1}p_{21} - \left(k_{+2} + k_{-1}\right)p_{11}, \\ \dfrac{dp_{00}}{dt} = 2\left(1 - p_{\text{n}}\right)k_{-1}p_{21} + k_{-1}p_{11}, \end{cases} \tag{S23}$$

where

$$k_{-1}(F) = k_{-1}^0 \exp\left(\frac{x_{-1}F}{k_{\text{B}}T}\right), \qquad k_2(F) = 2k_{-1}(F/2),$$

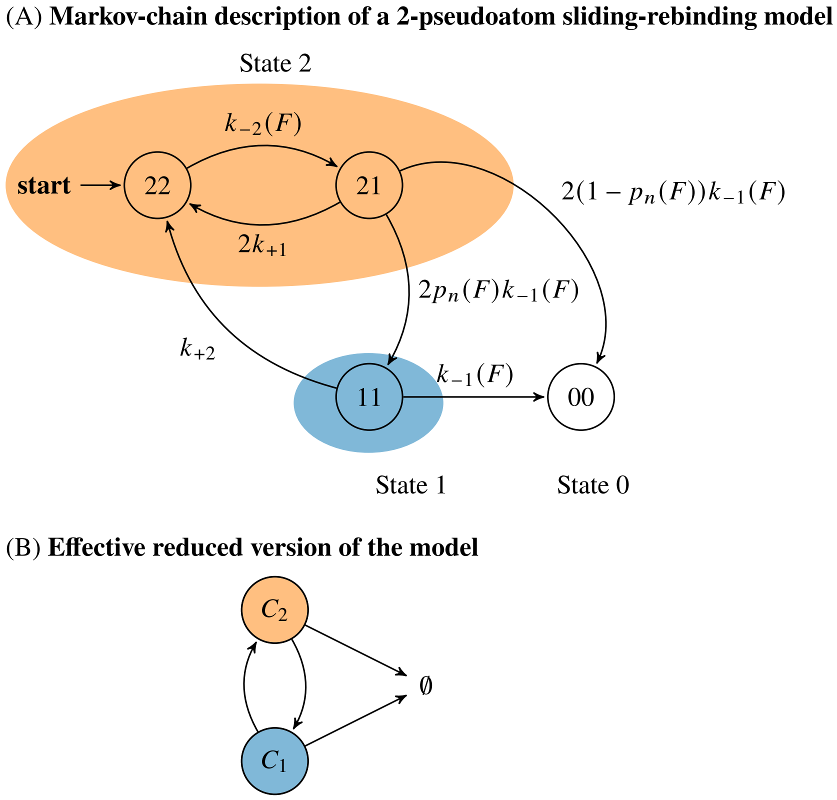


**Figure S7.** Markov-chain description of the sliding-rebinding model in Fig. S6. (A) The indices $(ij)$ label collective states, where $i$ is the maximum number of pseudoatom pairs and $j$ is the number of bound pairs. Transitions correspond to bond formation, bond breakage, and sliding with probability $p_n(F)$. The state (00) is the fully unbound state. (B) Effective reduced model obtained by grouping the colored highlighted states into two coarse-grained bound states, $C_2$ and $C_1$, and the unbound state $\emptyset$.

and the sliding-rebinding probability $p_n(F)$ was chosen in Ref. (15) to approximate the hinge-angle versus force curve obtained from molecular dynamics simulations.

This model naturally introduces multiple timescales (associated with the eigenvalues of the subsystem formed by $p_{22}$, $p_{21}$, and $p_{11}$), so the survival function $S(t) = 1 - p_{00}(t)$ generally exhibits several exponential components. For a generic SR model with $N$ pseudoatoms, one expects a wide variety of behaviors arising from the many underlying timescales. However, if the binding and rebinding rates of the pseudoatoms are comparable and there is a clear separation of timescales between fast internal relaxation and slow overall dissociation, the effective dynamics can be reduced in complexity (Fig. S7B), and as shown in the next subsection, the corresponding reduced model equations make this model mathematically equivalent to the TPSP model.

#### S2.6 Equivalence between TP, TSTP, BD, AD and SR models

The TSTP and AD models all introduce two dissociation pathways or bound states, differing in how they couple conformational changes to dissociation. In the AD model, the second state is force-stabilized; in the TSTP model, both states exist at zero force. Under Kramers' conditions (Appendix S1), these landscapes reduce to a Markov-chain description (18). A generic two-state scheme has the form of Fig. S3, where $C_0$ and $C_1$ are two bound states and $\emptyset$ is the unbound state. When transitions between $C_0$ and $C_1$ are much faster than unbinding, the bond survival probability takes a bi-exponential form,

$$S(t) = \omega e^{-k_1 t} + (1-\omega)e^{-k_2 t}, \tag{S24}$$

with two effective rates $k_1$, $k_2$ and a weight $\omega$. Thus, very different energy landscapes (TSTP, AD, BD, or SR) can yield indistinguishable survival curves once coarse-grained to this level, highlighting an intrinsic limitation of mechanism identification based solely on lifetime measurements.

Below, it is shown how the models TP, TSTP, AD, and SR are mathematically similar under suitable conditions, typically when fast molecular rearrangements are well separated in time from bond rupture.

**TP and AD models.** Pereverzev *et al.* (14) showed that the TP model is a limiting case of the allosteric (AD) model. Mathematically, this is the case when $k_1 \simeq k_2$, so $S(t) \sim e^{-k_1 t}$. Hereafter, we will not consider TP, as it cannot account for the experimental biphasic decay (we included it for historical reasons).

**TSTP and AD models.** Here we show how the AD model can be reinterpreted as a particular realization of the two-state, two-pathway scheme.

In the TSTP model, the bond survival probability $S(t)$ satisfies

$$\frac{dS(t)}{dt} = -k_{10}P_1(t) - k_{20}P_2(t), \tag{S25}$$

where $P_1(t)$ and $P_2(t)$ are the probabilities of occupying the bound states $C_1$ and $C_2$, and $k_{10}$, $k_{20}$ are the corresponding off-rates (see Eq. (S12)). In the allosteric model, the analogous survival probability $\tilde{S}(t)$ obeys,

$$\frac{d\tilde{S}(t)}{dt} = -k(t)\tilde{S}(t) = -\left[k_{10}Q_1(t) + k_{20}Q_2(t)\right] \cdot \tilde{S}(t), \tag{S26}$$

where $Q_1(t)$ and $Q_2(t)$ are the probabilities of being in the allosteric states $R_1L$ and $R_2L$ (Eq. (S16)), and $k(t)$ is the time-dependent effective off-rate defined in Eq. (S19).

By inspection of Eqs. (S25, S26), we can rewrite Eq. (S26) in terms of

$$\tilde{P}_1(t) = Q_1(t)\,\tilde{S}(t), \qquad \tilde{P}_2(t) = Q_2(t)\,\tilde{S}(t),$$

which play the role of populations of the two bound states in the allosteric picture. Then, Eq. (S26) becomes:

$$\frac{d\tilde{S}(t)}{dt} = -k_{10}\tilde{P}_1(t) - k_{20}\tilde{P}_2(t), \tag{S27}$$

which has the same structure as Eq. (S25). Considering that $Q_1(t) + Q_2(t) = 1$, the above definitions of $\tilde{P}_1(t)$ and $\tilde{P}_2(t)$ imply $\tilde{S}(t) = \tilde{P}_1(t) + \tilde{P}_2(t)$, $Q_1(t) = \tilde{P}_1(t)/(\tilde{P}_1(t) + \tilde{P}_2(t))$ and $Q_2(t) = \tilde{P}_2(t)/(\tilde{P}_1(t) + \tilde{P}_2(t))$. Therefore, differentiating $\tilde{P}_1(t)$ and $\tilde{P}_2(t)$ and substituting Eq. (S16) yields,

$$\begin{cases} \dfrac{d\tilde{P}_1(t)}{dt} &= -k_{12}\tilde{P}_1(t) + k_{21}\tilde{P}_2(t) - k(t)\,\tilde{P}_1(t), \\ \dfrac{d\tilde{P}_2(t)}{dt} &= k_{12}\tilde{P}_1(t) - k_{21}\tilde{P}_2(t) - k(t)\,\tilde{P}_2(t), \end{cases} \tag{S28}$$

where the effective rate $k(t)$ can be written as

$$k(t) = k_{10}Q_1(t) + k_{20}Q_2(t) = \frac{k_{10}\tilde{P}_1(t) + k_{20}\tilde{P}_2(t)}{\tilde{P}_1(t) + \tilde{P}_2(t)}. \tag{S29}$$

Comparing these equations with the master system (S12), we see that the allosteric model is mathematically equivalent to a two-state, two-pathway model in which both bound states $C_1$ and $C_2$ dissociate with the same instantaneous rate $k(t)$.

**BD and AD models.** The AD model has been shown to be a particular instance of the BD model (13). For completeness, we reproduce the derivation here. In BD, introduced in Ref. (12), the resulting allosteric contribution to the binding energy is

$$\Delta E_c(F) = (\Delta E_2 - \Delta E_1)\left[P_2(F) - P_2(0)\right],$$

where $\Delta E_1$ and $\Delta E_2$ denote the binding-potential depths in states 1 and 2. Combining this allosteric effect with the Bell lowering of the activation barrier yields

$$\Delta E(F) = \Delta E_0 + \Delta E_c(F) - Fx,$$

with $\Delta E_0 = (\Delta E_1 + \beta\Delta E_2)/(1+\beta)$ the zero-force barrier height. Since state 2 is much deeper than state 1 ($\Delta E_2 \gg \Delta E_1$), the allosteric energy change reduces to

$$\Delta E_c(F) = \frac{\Delta E_2 \beta\,[1-\exp(-2x_d F/k_B T)]}{\beta + \exp(-2x_d F/k_B T)}.$$

This expression predicts that the allosteric lowering of the potential well increases linearly at small forces and saturates at $\Delta E_2$ for $F \gg F_0$, as postulated for BD.

**TSTP and SR models.** In the previous subsection, we stated that the equations of the SR reduced model are also similar to those of the TSTP model. Here we show that this is the case.

For $N = 2$, grouping states according to the first index —as indicated by the colored regions in Fig. S7(A)— and assuming that states (22) and (21) are in quasi-steady state (21), we obtain an effective description in terms of two coarse-grained bound states, $C_2$ and $C_1$, and the unbound state $\emptyset$ (Fig. S7B). The corresponding equations are:

$$\begin{cases} \dfrac{dp_{(2)}}{dt} = k_{+2}p_{(1)} - \dfrac{2k_{-1}}{1+\frac{2k_{+1}}{k_{-2}}}\, p_{(2)}, \\ \dfrac{dp_{(1)}}{dt} = \dfrac{2p_{\mathrm{n}}k_{-1}}{1+\frac{2k_{+1}}{k_{-2}}}\, p_{(2)} - (k_{+2}+k_{-1})\, p_{(1)}, \\ \dfrac{dp_{\emptyset}}{dt} = -\dfrac{dS(t)}{dt} = \dfrac{2\,(1-p_{\mathrm{n}})\,k_{-1}}{1+\frac{2k_{+1}}{k_{-2}}}\, p_{(2)} + k_{-1}p_{(1)}, \end{cases} \tag{S30}$$

where we have used the stochastic quasi-steady state condition:

$$k_{-2}p_{22} = 2k_{+1}p_{21} \Rightarrow p_{(2)} = p_{21} + p_{22} = \left(1 + \frac{2k_{+1}}{k_{-2}}\right)p_{21} = \left(1 + \frac{k_{-2}}{2k_{+1}}\right)p_{22}. \tag{S31}$$

Thus, despite its structural richness, the SR model can effectively collapse onto a simpler kinetic scheme at the level of experimentally accessible observables, illustrating the problem of the broader issue of mechanistic non-uniqueness in catch-bond modeling (6, 39). However, the effective rates in the SR reduction (Eq. S30) involve *ratios* of force-dependent Bell rates, *e.g.*, $k_{\mathrm{eff}} \propto k_{-1}k_{-2}/(k_{+1}+k_{-2})$, which produces non-Bell-like force dependence. This provides a practical experimental signature for this model: measuring the force dependence of effective rates with sufficient precision can discriminate sliding-rebinding mechanisms from allosteric or two-pathway schemes.

As discussed in the main text, the general solution of the master equation (S31) can be expressed as the sum of exponentials (see Eq. (8) in Sec . 2, so we omit the explicit derivation for the SR model here, given that their master equations are equivalent as shown above.

#### S3 CFPR MODEL: DERIVATION OF THE MEAN LIFETIME FOR THE BIRTH–DEATH LADDER

This appendix derives the mean absorption time used in the CFPR model.

##### S3.1 Mean-field approximation (closed form)

Consider a birth–death process on states $i \in \{0,\dots,N\}$ with $i = 0$ absorbing. Under the *mean-field* simplification, the per-state death rate $\mu(F)$ is taken as constant (independent of $i$), whereas the full load-sharing model uses the state-dependent rate $i\,\mu(F/i)$ (main text, Section 4). With constant rates, the backward equation for the mean absorption time $\tau_m$ is

$$(b+\mu)\tau_i - b\,\tau_{i+1} - \mu\,\tau_{i-1} = 1, \quad 1 \le i \le N-1,$$

whose solution is Eq. (16) in the main text, namely:

$$\tau_m = \frac{m}{\mu(F) - b(F)}.$$

Hence, catch-like behavior arises when the net drift $\mu(F) - b(F)$ is non-monotone in $F$.

#### S3.2 Exact MFPT from state 1 (load-sharing model)

For the full load-sharing model, death rates are state-dependent: $d_{i,i-1}(F) = i\,\mu_0\, e^{\Delta x F/(i\,k_B T)}$. Using the standard linear-chain absorption-time formula (24), the mean first-passage time from state 1 to the absorbing state 0 is

$$\tau_1(F) = \frac{1}{\mu_0\, e^{\Delta x F/k_B T}} + \sum_{i=2}^{N} \frac{P_i(F)}{i!\; \mu_0^i\; e^{H_i\, \Delta x F/k_B T}}, \tag{S32}$$

where $H_i = \sum_{k=1}^{i} 1/k$ is the $i$-th harmonic number, and

$$P_i(F) = \prod_{j=1}^{i-1} \left(\lambda(F) + k_{\text{on}}^0 (N-j)\, e^{-\Delta x_{\text{on}} F/k_B T}\right)$$

is the product of birth rates at states $1, \ldots, i-1$. The harmonic numbers appear in the denominator because load-sharing gives $\prod_{j=1}^{i} d_{j,j-1} = i!\, \mu_0^i\, e^{H_i \Delta x F/k_B T}$, which follows from $\sum_{j=1}^{i} 1/j = H_i$.

For $F \longrightarrow \infty$, the exponential in the death rates (denominator) dominates, so $\tau_1(F) \longrightarrow 0$. So, a necessary condition for catch behavior is that $d\tau_1/dF|_{F=0} > 0$ (this applies also to the general case, $\tau_m$, where initially one has $m$ engaged RBCs). This can be visually seen in Fig. S8, which shows $\tau_m(F)$ and its derivative for three values of $\lambda_1$ (the force-dependent part of the birth rate). For $\lambda_1 = 0$ (pure slip), $d\tau_m/dF < 0$ everywhere. Increasing $\lambda_1$ generates a catch window where $d\tau_m/dF > 0$. The catch–slip transition occurs at the force where $d\tau_m/dF = 0$ (horizontal dashed line in the bottom panel).

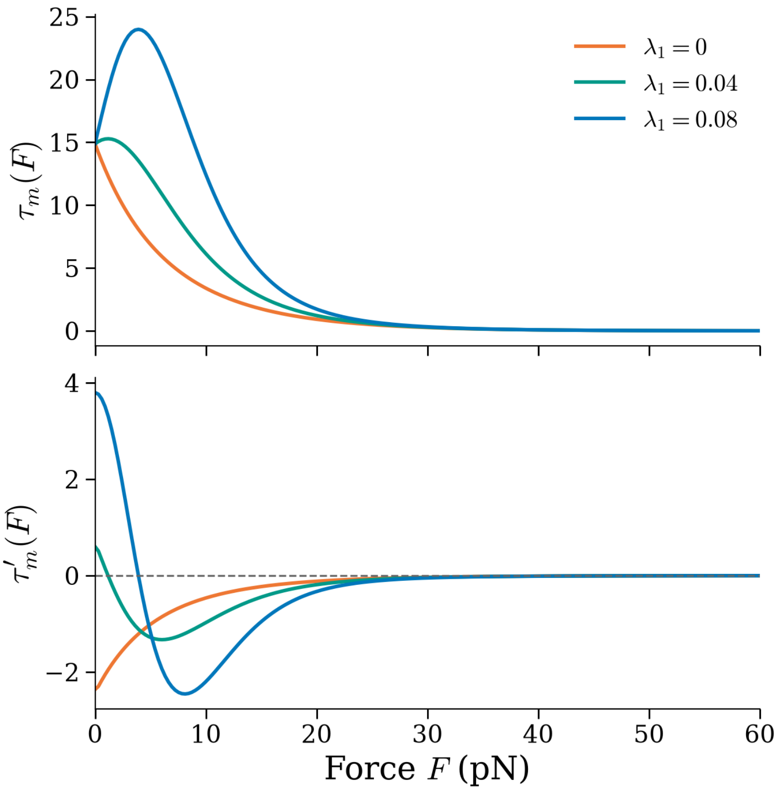


**Figure S8. Mean interphase lifetime $\tau_m(F)$ and its force derivative for three values of $\lambda_1$.** *Top:* $\tau_m(F)$ vs. applied force. *Bottom:* $d\tau_m/dF$ vs. force; horizontal dashed line at zero marks the catch–slip transition. For $\lambda_1 = 0$ (pure slip), $d\tau_m/dF < 0$ everywhere. Increasing $\lambda_1$ generates a catch window where $d\tau_m/dF > 0$. Fixed parameters as in Table S1 (catch regime).

**Catch-bond threshold (no rebinding, $\lambda_0 = 0$).** Setting $k_{\text{on}}^0 \rightarrow 0$ and $\lambda(F) = \lambda_1 F$ simplifies $P_i(F) = (\lambda_1 F)^{i-1}$. Differentiating $\tau_1$ with respect to $F$ and evaluating at $F = 0$ (using $H_1 = 1, T_1'(0) = -\mu_0^{-1}\Delta x/k_B T$, and $T_n'(0) = 0$ for $n \geq 3$ when $\lambda_0 = 0$) gives

$$\left.\frac{d\tau_1}{dF}\right|_{F=0} = -\frac{\Delta x}{k_B T\,\mu_0} + \frac{\lambda_1}{2\mu_0^2}.$$

The factor of 2 in the second term arises from the $i! = 2! = 2$ factorial in the $i = 2$ term of Eq. (S32): differentiating $P_2/(2!\,\mu_0^2) = \lambda_1 F/(2\mu_0^2)$ at $F = 0$ yields $\lambda_1/(2\mu_0^2)$. When $\lambda_0 = 0$, all $i \geq 3$ terms vanish at $F = 0$ since $P_i \propto (\lambda_1 F)^{i-1}$, so only the $i = 1$ (single-bond escape) and $i = 2$ (first load-sharing correction) terms contribute. The catch condition $d\tau_1/dF|_{F=0} > 0$ therefore requires

$$\lambda_1 > \frac{2\mu_0\,\Delta x}{k_B T} = \frac{2\mu_0}{F_b}, \tag{S33}$$

where $F_b = k_B T/\Delta x$. This is *strictly stronger* than the mean-field condition $\lambda_1 > \mu_0/F_b$; both are satisfied for our parameters ($\lambda_1 = 0.08\,\text{s}^{-1}\text{pN}^{-1}$, $\mu_0/F_b \approx 0.012\,\text{s}^{-1}\text{pN}^{-1}$, $2\mu_0/F_b \approx 0.024\,\text{s}^{-1}\text{pN}^{-1}$).

**General threshold $T_N(\alpha)$.** For general $N$ and $\lambda_0 \geq 0$ (still without rebinding, $k_{\text{on}}^0 = 0$), we set $\alpha = \lambda_0/\mu_0$ and differentiate $\tau_1(F)$ at $F = 0$ using $\lambda(F) = \lambda_0 + \lambda_1 F$ and $P_i(F) = (\lambda_0 + \lambda_1 F)^{i-1}$:

$$\left.\frac{d\tau_1}{dF}\right|_{F=0} = -\frac{1}{F_b\mu_0}\left[1 + \sum_{i=2}^{N} \frac{H_i\,\alpha^{i-1}}{i!}\right] + \frac{\lambda_1}{\mu_0^2}\sum_{i=2}^{N}\frac{(i-1)\,\alpha^{i-2}}{i!}.$$

The catch condition $d\tau_1/dF|_{F=0} > 0$ gives $\lambda_1 > T_N(\alpha)\cdot(\mu_0/F_b)$ with

$$T_N(\alpha) = \frac{1 + \displaystyle\sum_{n=2}^{N}\frac{H_n\,\alpha^{n-1}}{n!}}{\displaystyle\sum_{n=2}^{N}\frac{(n-1)\,\alpha^{n-2}}{n!}}, \tag{S34}$$

where $H_n = \sum_{k=1}^{n} 1/k$ is the $n$-th harmonic number. Setting $\alpha = 0$: the numerator is 1 (all $\alpha^{n-1}$ terms vanish for $n \geq 2$) and the denominator reduces to $(2-1)/2! = 1/2$ (only the $n = 2$ term survives since $0^{n-2} = 0$ for $n \geq 3$), giving $T_N(0) = 2$ for all $N \geq 2$, consistent with Eq. (S33).

### S4 CFPR MODEL: SYNTHETIC TRAJECTORY GENERATION AND BAYESIAN INFERENCE

#### S4.1 Synthetic trajectory generation

Here, we implement the full model described in Fig. 2 to test whether it can extract information from Molecular Dynamics (MD) trajectories. Because the adapted Classical Trajectory Monte Carlo (CTMC) method assumes a constant applied force, the corresponding MD protocol should be quasistatic: apply a fixed strain, allow the system to mechanically equilibrate, then record RBC engagement and disengagement events until the complex reaches the fully unbound (absorbing) state (29). Synthetic contact-count trajectories $n(t) \in \{0, \ldots, N\}$ were generated via the Gillespie algorithm (44) under a CTMC of the CFPR model (Section 4). At each step, death rate $d(n, F) = n\,k_{\text{off}}^0 \exp(F\Delta x/(nk_{\text{B}}T))$ and birth rate $b_{n,n+1}(F) = r_{n,n+1}(F) + \lambda(F)$ (Eqs. 11–20) were computed; holding time $\Delta t \sim \text{Exponential}(d + b)$ was drawn; event type was chosen proportionally to the corresponding rate; state $n$ was updated. Simulations terminated at absorption ($n = 0$) or $t_{\max} = 200$ s (censored; > 99% of trajectories reached absorption). Two parameter regimes (Table S1) were used: *slip* ($\lambda_1 = 0$, $\lambda_2 = 0.1$ pN$^{-1}$) and *catch* ($\lambda_1 = 0.08$ s$^{-1}$ pN$^{-1}$, $\lambda_2 = 0.08$ pN$^{-1}$). Force values were $F \in \{0, 2, 5, 10, 15, 20, 30, 40, 60\}$ pN, and 200 replicates were performed per force.

The parameter values in Table S1 are anchored to the receptor–ligand and TCR–pMHC biophysics literature; those that cannot be measured directly are chosen only to place the model in a physically plausible regime, to be constrained later by molecular dynamics or single-molecule data. The intrinsic zero-force off-rate $k_{\text{off}}^0 \approx 0.1$ s$^{-1}$ (a bound lifetime of 10 s) lies within the range of surface-plasmon-resonance dissociation rates of strong TCR–pMHC agonists ($\sim$ 0.01–0.5 s$^{-1}$) (45); note that two-dimensional force-clamp assays report faster intrinsic off-rates ($\sim$ 1–10 s$^{-1}$) (7, 34), so $k_{\text{off}}^0$ should be read as a solution-phase value.

The transition-state distance $\Delta x = 0.5$ nm coincides with the canonical outer-barrier compliance of the streptavidin–biotin bond and lies mid-range for non-covalent protein–ligand slip barriers (0.1–1 nm) (1, 46). The valency $N = 5$ is the coarse-grained number of load-bearing non-covalent residue-binding contacts (RBCs) at the interface, consistent with the small set ($\sim$ 5–10) of hydrogen bonds and the $\sim$ 3–6 energetically dominant "hotspot" contacts resolved in TCR–pMHC structures and alanine-scanning studies (29, 47, 48). The per-contact rebinding rate $k^0_{\mathrm{on}} \approx 0.02\ \mathrm{s}^{-1}$ is obtained from the two-dimensional agonist on-rate of Ref. (35) ($A_c k_{\mathrm{on}} \approx 1.2 \times 10^{-2}\ \mu\mathrm{m}^4\ \mathrm{s}^{-1}$) divided by a contact area $A_c \approx 1\ \mu\mathrm{m}^2$, giving $\approx 0.012\ \mathrm{s}^{-1}$; the rebinding distance $\Delta x_{\mathrm{on}} \approx 0.2$ nm has the sign and order of magnitude implied by the force-suppression of association (49). As shown in Section 6 of the main text, $k^0_{\mathrm{on}}$ and $\Delta x_{\mathrm{on}}$ are structurally non-identifiable parameters when only force–lifetime data are used, so these order-of-magnitude estimates are the appropriate level of commitment.

The engagement rate $\lambda(F) = \lambda_1 F\, e^{-\lambda_2 F}$ (we deliberately set the force-free rebinding baseline $\lambda_0 = 0$, the most conservative choice, so that engagement is entirely force-driven) encodes mechano-nucleation: lateral load recruits new potential molecular contacts, motivated by molecular dynamics showing that pulling exposes an otherwise cryptic binding pocket (29). No direct measurement of a TCR force-induced nucleation slope exists, so $\lambda_1 \approx 0.08\ \mathrm{s}^{-1}\mathrm{pN}^{-1}$ is a proof-of-concept value; its sign and existence are supported by direct observations that force switches receptor–ligand complexes to faster-on-rate states near 10 pN (11–17-fold for the von Willebrand factor A1/GPIb$\alpha$ complex (50)) and by the more general phenomenon of force exposing cryptic binding sites (51). The high-force suppression $\lambda_2 \approx 0.08\ \mathrm{pN}^{-1}$ is not arbitrary: read as a Bell transition-state distance $\lambda_2 = x_\beta / k_B T$, it gives $x_\beta \approx 0.33$ nm, a standard protein-bond reaction coordinate, and sets a characteristic force $1/\lambda_2 = 12.5$ pN. Consequently, the emergent CFPR catches maximum falls at $F^* \approx$ 5–15 pN — in agreement, without any fitting, with the measured TCR–pMHC catch-bond peak forces of $\sim$ 10 pN (7), $\sim$ 15 pN (34), and $\sim$ 10–16 pN (52).

**Table S1.** Simulation parameter sets.

| Parameter | Slip | Catch |
|---|---|---|
| $N$ | 5 | 5 |
| $k^0_{\mathrm{off}}$ ($\mathrm{s}^{-1}$) | 0.10 | 0.10 |
| $\Delta x$ (nm) | 0.50 | 0.50 |
| $k^0_{\mathrm{on}}$ ($\mathrm{s}^{-1}$) | 0.02 | 0.02 |
| $\Delta x_{\mathrm{on}}$ (nm) | 0.20 | 0.20 |
| $\lambda_0$ | 0.0 | 0.0 |
| $\lambda_1$ ($\mathrm{s}^{-1}\ \mathrm{pN}^{-1}$) | 0.0 | 0.08 |
| $\lambda_2$ ($\mathrm{pN}^{-1}$) | 0.1 | 0.08 |

#### S4.2 Force–lifetime summary

Mean lifetimes were estimated as the restricted mean survival time (RMST) of the Kaplan–Meier curve (53), integrated from $t = 0$ to $t_{\mathrm{max}}$ via a left Riemann sum over the Kaplan–Meier step function. This estimator is exact for right-continuous step functions and accounts for right-censoring.

#### S4.3 Bayesian parameter inference

The 200 trajectories per force generated in Section S4.1 served two separate purposes: all 200 were used for the Kaplan–Meier force–lifetime summary (Section S4.2), while a random subsample of 10 per force was drawn independently for Bayesian inference (totalling 90 trajectories; the full CTMC path likelihood requires evaluating transition rates for each interval row, making the larger dataset computationally prohibitive for 4-chain Hamiltonian Monte-Carlo (HMC) using the standard diagnostic tests (54)). Parameters $k^0_{\mathrm{off}}, \Delta x, k^0_{\mathrm{on}}, \Delta x_{\mathrm{on}}, \lambda_0, \lambda_1, \lambda_2$, and $N$ were inferred from these 10 trajectories per force value (1266 interval rows) using the exact CTMC path log-likelihood:

$$\log \mathcal{L}(\boldsymbol{\theta}) = \sum_{k=1}^{J} \log W_{n_{k-1},n_k}(F(t_k)) - \int_0^{\tau} \sum_{j \neq n(s)} W_{n(s),j}(F(s))\, ds, \tag{S35}$$

where $J$ is the number of transitions, $t_k$ are jump times, and $W_{u,v}$ are transition intensities. Sampling used PyMC (55): No-U-Turn Sampler (NUTS) (56) for seven continuous parameters and Categorical Gibbs–Metropolis for discrete $N \in \{1, \dots, 10\}$ with Poisson($\mu = 5$) prior. Four chains, 2000 draws each, 1000 tuning steps; $\hat{R} < 1.01$ for all parameters; 1 divergent transition in 8000 draws. Exact recovery of $N = 5$ across all four chains (zero between-chain disagreement) confirms adequate mixing of the discrete component. Wall time: 24 s. Priors used for each parameter are indicated in Table S2.

**Table S2.** Prior distributions for Bayesian inference.

| Parameter | Prior | Hyperparameters | Support |
|---|---|---|---|
| $k^0_{\text{off}}$ | LogNormal | $\mu_{\ln} = -2.3, \sigma = 1.0$ | $(0, \infty)$ |
| $\Delta x$ | Uniform | | $[0.01, 2.0]$ |
| $k^0_{\text{on}}$ | LogNormal | $\mu_{\ln} = -4.0, \sigma = 1.5$ | $(0, \infty)$ |
| $\Delta x_{\text{on}}$ | Uniform | | $[0.01, 2.0]$ |
| $\lambda_0$ | HalfNormal | $\sigma = 0.5$ | $[0, \infty)$ |
| $\lambda_1$ | HalfNormal | $\sigma = 0.5$ | $[0, \infty)$ |
| $\lambda_2$ | Uniform | | $[0.001, 1.0]$ |
| $N$ | Categorical (Poisson-truncated) | $\mu = 5$ | $\{1, \dots, 10\}$ |

##### S4.4 Posterior predictive checks

100 posterior samples were drawn and, for each, 20 Gillespie trajectories were simulated per force value. The 5th–95th percentile of Kaplan–Meier mean bond lifetimes across posterior samples defines a 90% posterior predictive credible band. PPC coverage is the fraction of observed mean lifetimes inside this band.

#### S5 CFPR MODEL: SENSITIVITY ANALYSIS OF CFPR ENGAGEMENT RATE FUNCTIONAL FORM

The proof-of-concept CFPR model in the main text uses a linear engagement rate $\lambda(F) = \lambda_0 + \lambda_1 F$. Here, we test the robustness of the qualitative bifurcation behavior to alternative functional forms.

We consider three alternative forms motivated by different physical mechanisms:

**(1) Exponential engagement:** $\lambda(F) = \lambda_0 \exp(\alpha F)$, representing cooperative bond formation with a characteristic force scale $1/\alpha$.

**(2) Saturating engagement:** $\lambda(F) = \lambda_0 + \lambda_{\max}[1 - \exp(-bF)]$, representing a finite pool of recruitable bonds that saturates at high forces.

**(3) Geometry-dependent engagement:** $\lambda(F) = \lambda_0[1 + \cos(\phi(F))]$, where $\phi(F) = \phi_0 \exp(-F/F_c)$ models pulling-angle-dependent molecular alignment.

For each form, we compute the mean lifetime $\tau_m(F)$ using Eq. (16) and identify the location of the catch-to-slip crossover $F^*$. Results show that:

- All three forms produce qualitatively similar slip→catch→slip crossover behavior when parameters are tuned to satisfy the bias condition $\mu(F) > \lambda(F)$.
- The quantitative location of $F^*$ varies by ±30% relative to the primary suppressed-form value $F^* \approx 6.1$ pN (Table 2 parameters), indicating that precise predictions require molecular-scale constraints.
- The exponential form yields the sharpest crossover, while the saturating form produces a broader catch regime, consistent with intuition about limited bond recruitment.

These results confirm that the CFPR mechanism—emergent catch behavior from force-driven synapse reorganization—is robust to the specific parameterization of $\lambda(F)$, though quantitative fitting to experimental data will require constraining this function via molecular dynamics or structural measurements.

#### S6 CFPR MODEL: BAYESIAN INFERENCE DIAGNOSTICS AND SUPPLEMENTARY ANALYSES

This appendix presents supplementary analyses supporting the results in Section 6: posterior marginal distributions, convergence diagnostics, pairwise parameter correlations, predictive checks, and robustness tests.

##### S6.1 Posterior marginal distributions and convergence

Trace plots and marginal posterior histograms for all eight CFPR parameters (Fig. S9 and Fig. S10) were obtained from four Hamiltonian Monte Carlo chains (2000 draws each, 1000 tuning steps). Convergence diagnostics: $\hat{R} < 1.01$ for all parameters, 1 divergent transition in 8000 total draws.

Table S3 lists posterior means, 95% credible intervals, and true values for the catch parameter regime.

**Table S3.** Posterior summary for CFPR parameters.

| Parameter | True | Mean | SD | 95% CI | Recovery |
|---|---|---|---|---|---|
| $k^0_{\mathrm{off}}$ ($\mathrm{s}^{-1}$) | 0.100 | 0.096 | 0.004 | [0.088, 0.104] | Good |
| $\Delta x$ (nm) | 0.500 | 0.515 | 0.015 | [0.484, 0.544] | Good |
| $k^0_{\mathrm{on}}$ ($\mathrm{s}^{-1}$) | 0.020 | 0.008 | 0.005 | [0.001, 0.019] | Poor |
| $\Delta x_{\mathrm{on}}$ (nm) | 0.200 | 1.003 | 0.568 | [0.064, 1.952] | Not identified |
| $\lambda_0$ | 0.0 | 0.023 | 0.015 | [0.001, 0.057] | Fair |
| $\lambda_1$ ($\mathrm{s}^{-1}\,\mathrm{pN}^{-1}$) | 0.080 | 0.100 | 0.010 | [0.080, 0.121] | Good |
| $\lambda_2$ ($\mathrm{pN}^{-1}$) | 0.080 | 0.092 | 0.008 | [0.076, 0.109] | Good |
| $N$ | 5 | 5.0 | 0.0 | [5, 5] | Exact |

The dissociation arm ($k^0_{\mathrm{off}}$, $\Delta x$) and immigration arm ($\lambda_1$, $\lambda_2$, $N$) are well identified. Posterior means for $k^0_{\mathrm{off}}$ and $\Delta x$ recover true values within 4% and 3% respectively; $\lambda_1$ and $\lambda_2$ show moderate finite-data bias (25% and 15% respectively) but true values fall within their 95% credible intervals; and $N = 5$ is recovered exactly. The standard-rebinding pair ($k^0_{\mathrm{on}}$, $\Delta x_{\mathrm{on}}$) is not identified: the posterior for $\Delta x_{\mathrm{on}}$ is nearly flat across its prior support $[0.01, 2.0]$ nm, and the 95% CI for $k^0_{\mathrm{on}}$ is $[0.001, 0.019]\,\mathrm{s}^{-1}$ with the true value $0.020\,\mathrm{s}^{-1}$ lying just above the upper bound — a formal coverage failure for this parameter (the true value falls at the 98.3rd posterior percentile, inside the 99% CI $[0.001, 0.021]\,\mathrm{s}^{-1}$). This outcome is consistent with structural non-identifiability: the posterior mass is concentrated below the true value because force-induced immigration dominates the birth rate across the inference force range, leaving rebinding weakly constrained by the data (see Fig. S11).

##### S6.2 Pairwise parameter correlations

The complete joint posterior is shown in Fig. S11, which displays all 21 pairwise marginals together with the seven one-dimensional marginals. Three structural features stand out. The immigration pair $(\lambda_1, \lambda_2)$ is positively correlated ($r \approx +0.82$): a steeper linear rise is partially offset by faster suppression at high force. The off-rate pair $(k^0_{\mathrm{off}}, \Delta x)$ is moderately anti-correlated ($r \approx -0.55$): a larger Bell distance compensates a smaller prefactor. The standard-rebinding pair $(k^0_{\mathrm{on}}, \Delta x_{\mathrm{on}})$ is statistically independent ($r \approx +0.03$), with a flat $\Delta x_{\mathrm{on}}$ marginal spanning its full prior support and no compensatory ridge — the structural signature of non-identifiability, since force-induced immigration dominates the birth rate across the inference force range and leaves no information to constrain rebinding.

##### S6.3 Posterior predictive checks

Figure 4b (main text) shows that the inferred model reproduces the observed mean lifetimes across the full force range. To resolve the full predictive lifetime *distribution* at selected forces rather than only its mean, we drew a much larger predictive ensemble — 300 posterior samples with 60 Gillespie replicates per force ($\approx 1.8 \times 10^4$ predictive lifetimes per force value, two orders of magnitude above the band computation above). The resulting densities (Fig. S12) are essentially free of Monte-Carlo noise and reproduce the observed lifetime distributions across the force range, including the long right tail at low force and the sharp compression at high force. The agreement of the entire distribution shape — not just the mean — indicates

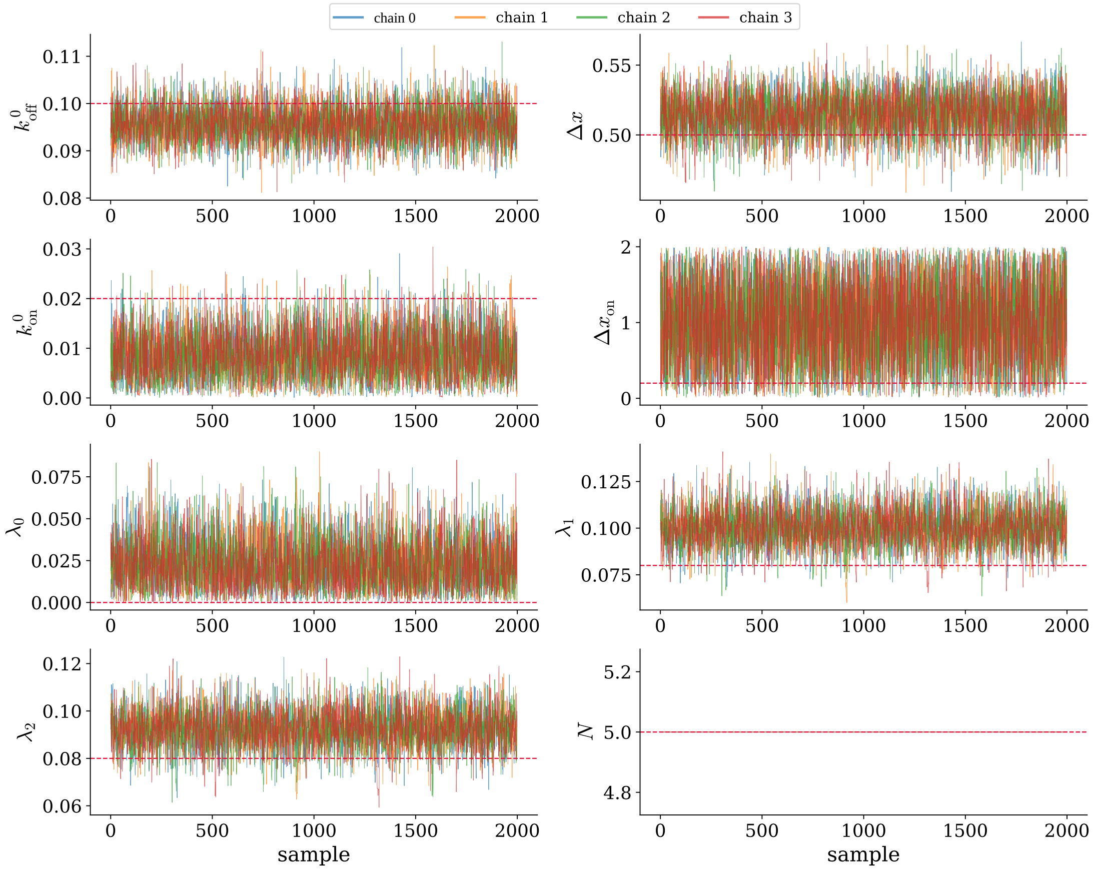


**Figure S9. Trace plots for CFPR parameters.** Four chains (colored) show good mixing for $k^0_{\text{off}}$, $\Delta x$, $\lambda_1$, $\lambda_2$, and $N$. Notably, the number of RBCs is identified without any uncertainty. In contrast, the simulated data contain insufficient information about $\Delta x_{\text{on}}$ to reconstruct this parameter. The horizontal line is the actual value used to generate the synthetic data.

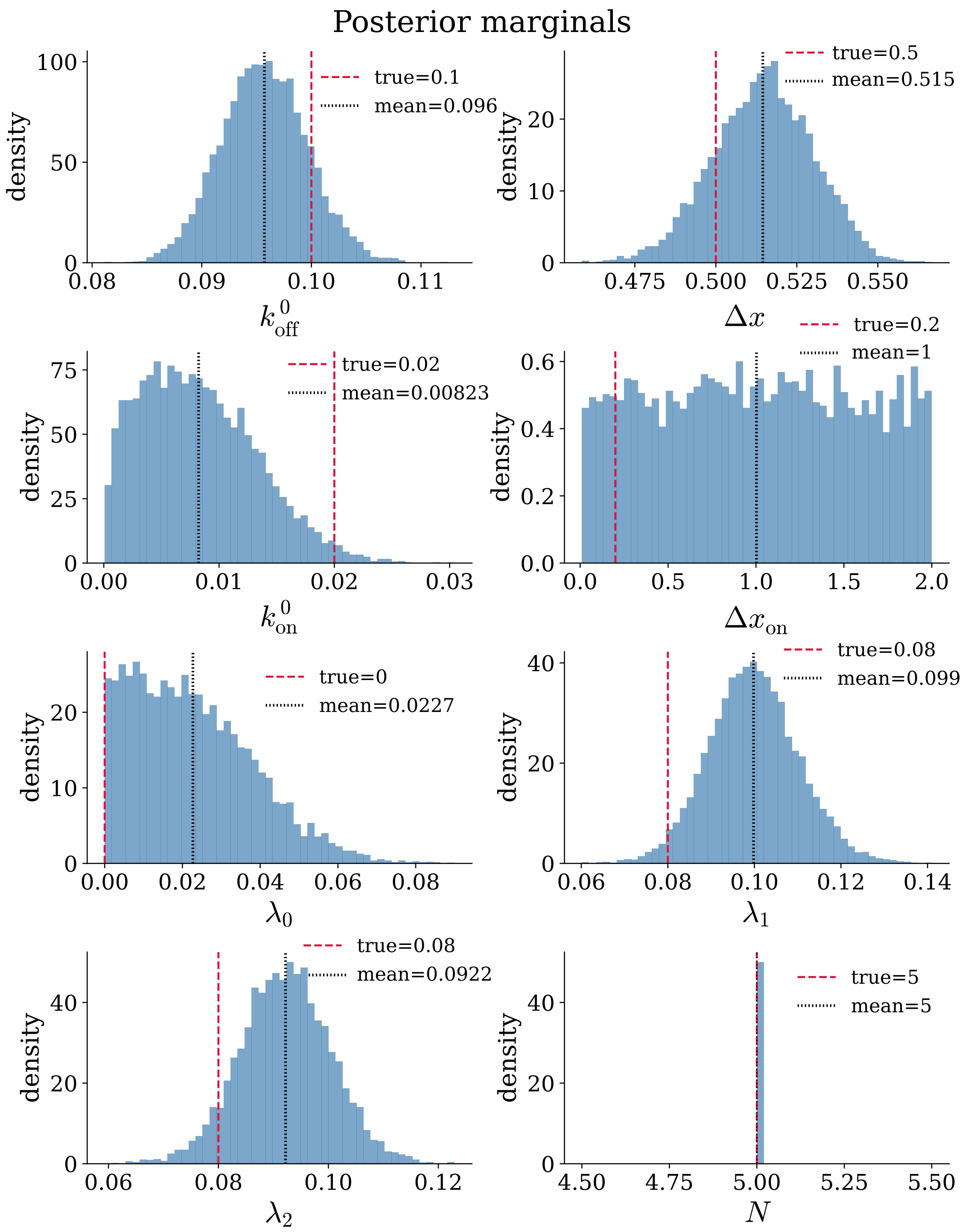


**Figure S10. Marginal posterior distributions.** Histograms (blue) with true parameter values (red dashed lines). The dissociation and immigration parameters are sharply peaked around their true values, whereas $\lambda_0$, $k^0_{\mathrm{on}}$, and $\Delta x_{\mathrm{on}}$ are widely spread, and the first two are highly correlated.

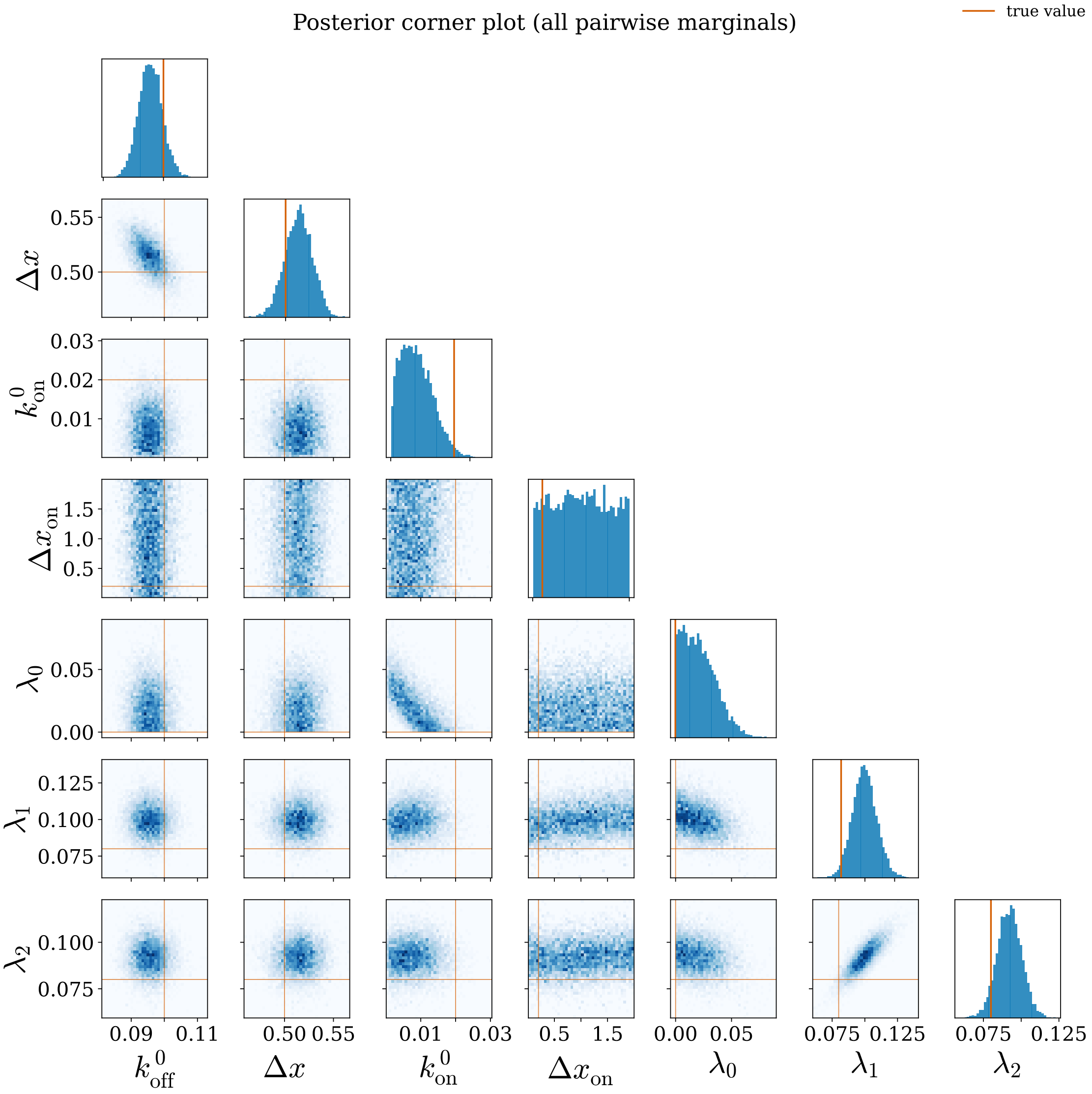


**Figure S11. Full posterior pair plots.** All 21 pairwise joint densities (lower triangle) and the seven one-dimensional marginals (diagonal) for the continuous CFPR parameters; red lines mark the data-generating values. The dissociation $(k_{\mathrm{off}}^{0}, \Delta x)$ and immigration $(\lambda_1, \lambda_2)$ compensatory ridges are visible, whereas the rebinding pair $(k_{\mathrm{on}}^{0}, \Delta x_{\mathrm{on}})$ shows a flat $\Delta x_{\mathrm{on}}$ marginal and no joint constraint.

that the discrepancies at the level of $\tau(F)$ reflect finite-data posterior bias rather than structural model misfit.

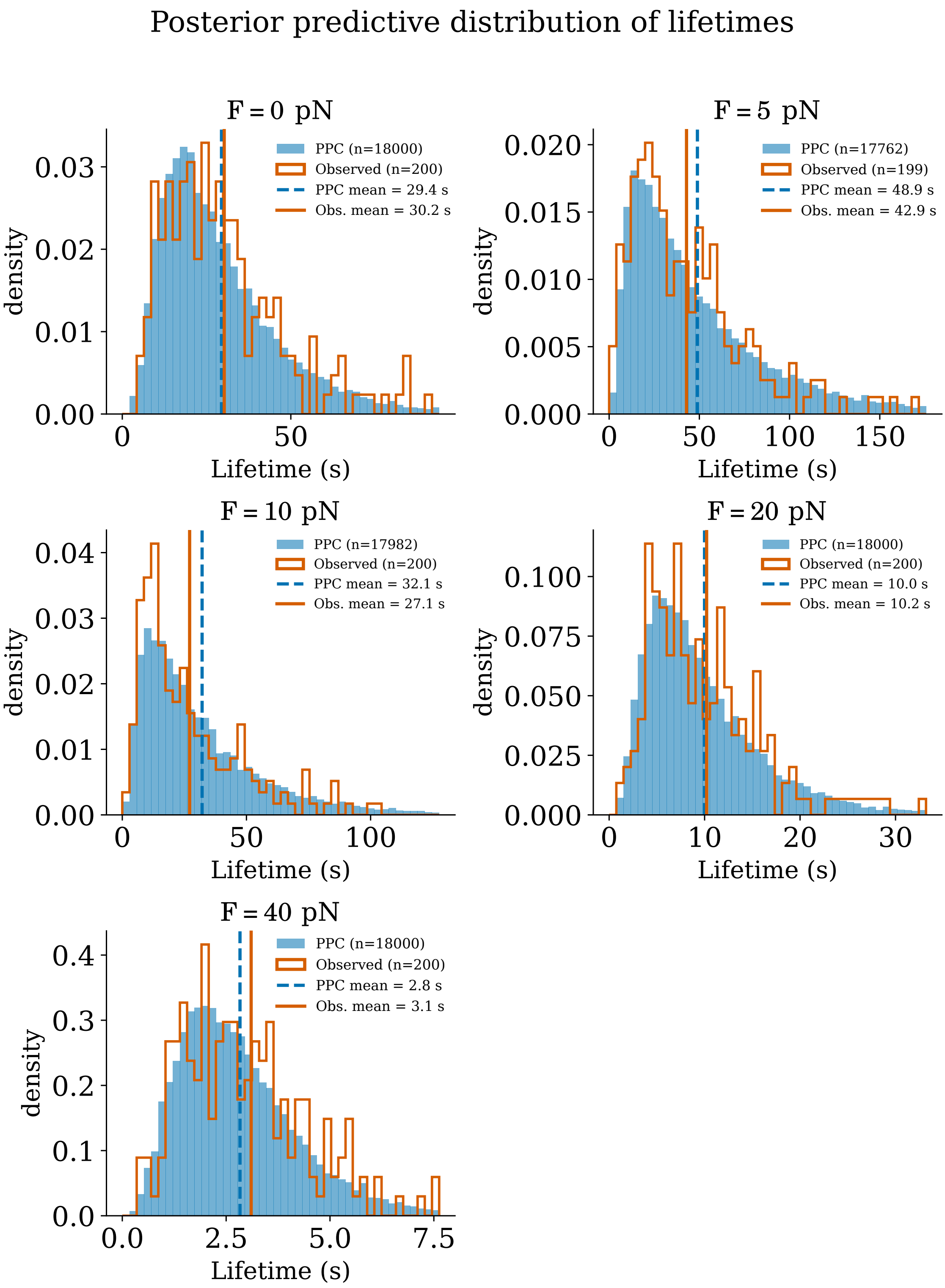


**Figure S12. Posterior predictive lifetime distributions.** Pooled posterior predictive lifetimes (filled blue, $n \approx 1.8 \times 10^4$ per force; 300 posterior draws $\times$ 60 replicates) versus observed lifetimes (red step, $n = 200$) at $F = 0, 5, 10, 20, 40$ pN. Vertical lines mark the posterior predictive mean (dashed blue) and the observed mean (solid red) lifetime at each force. The large ensemble resolves the full distribution, confirming that the fitted model captures both the location (mean) and the shape of the lifetime distribution at every force.

**Table S4. Per-force posterior predictive coverage summary.** Observed KM-RMST vs. 90% posterior predictive band ($q_5$–$q_{95}$ across 100 posterior samples × 20 replicates per force). All nine values lie inside the band (PPC coverage = 9/9). Bayesian predictive $p$-values (fraction of PPC samples with mean lifetime ≥ observed) range from 0.26 to 0.82; see main text Section S4.4.

| $F$ (pN) | Observed (s) | PPC mean (s) | 90% CI (s) | Inside |
|---|---|---|---|---|
| 0 | 30.24 | 30.19 | 24.4–38.6 | ✓ |
| 2 | 37.54 | 44.92 | 31.4–58.3 | ✓ |
| 5 | 43.68 | 53.78 | 37.9–72.7 | ✓ |
| 10 | 27.12 | 33.09 | 24.3–44.3 | ✓ |
| 15 | 16.77 | 17.03 | 13.4–20.9 | ✓ |
| 20 | 10.21 | 10.30 | 8.2–12.9 | ✓ |
| 30 | 5.19 | 4.98 | 4.0–6.1 | ✓ |
| 40 | 3.10 | 2.89 | 2.2–3.6 | ✓ |
| 60 | 1.25 | 1.34 | 1.1–1.6 | ✓ |

#### S6.4 Effective-rate non-linearity for sliding–rebinding

The reduced SR model (Appendix S2, Eq. S30) yields effective dissociation rates of the form

$$k_{\text{eff}}(F) = \frac{2k_{-1}(F)}{1 + 2k_{+1}(F)\,/\,k_{-2}(F)},$$

where each microscopic rate follows the Bell form: $k_{\pm i}(F) = k^0_{\pm i}\exp(x_{\pm i}F/k_BT)$. Substituting gives

$$k_{\text{eff}}(F) = \frac{2k^0_{-1}e^{x_{-1}F/k_BT}}{1 + \frac{2k^0_{+1}}{k^0_{-2}}e^{(x_{+1}-x_{-2})F/k_BT}}.$$

The denominator contains a ratio of exponentials, so $\ln k_{\text{eff}}(F)$ is not linear in $F$ (unlike a pure Bell rate). For $x_{+1} < x_{-2}$, the denominator grows with force, causing $\ln k_{\text{eff}}(F)$ to bend downward at high forces, a qualitative signature that distinguishes SR from TSTP and AD mechanisms (which produce purely exponential effective rates under timescale separation). This curvature over the 0–30 pN range provides an experimental signature distinguishing SR from TSTP and AD mechanisms.

### S7 CFPR MODEL PHASE DIAGRAMS

#### Model and phase criterion

The CFPR CTMC (Section 4) has death and birth rates

$$d(n,F) = n\,k^0_{\text{off}}\exp\left(\frac{F\,\Delta x}{n\,k_BT}\right), \tag{S36}$$

$$b(n,F) = (N-n)\,k^0_{\text{on}}\exp\left(-\frac{F\,\Delta x_{\text{on}}}{k_BT}\right) + (\lambda_0 + \lambda_1 F)\,e^{-\lambda_2 F}. \tag{S37}$$

The mean interphase lifetime $\tau_m(F)$ is computed numerically from the full absorbing CTMC by exact MFPT (matrix inversion). The catch/slip boundary is defined by the sign of

$$\left.\frac{d\tau_m}{dF}\right|_{F=0}: \quad \blacksquare\ \text{catch regime } (>0), \quad \blacksquare\ \text{slip regime } (\le 0).$$

Each phase diagram sweeps two parameters over the ranges in Table S5 while holding remaining parameters at the catch-regime defaults (Table S1): $k^0_{\text{off}} = 0.1\ \text{s}^{-1}$, $\Delta x = 0.5$ nm, $k^0_{\text{on}} = 0.02\ \text{s}^{-1}$, $\Delta x_{\text{on}} = 0.2$ nm, $\lambda_0 = 0$, $\lambda_1 = 0.08\ \text{s}^{-1}\,\text{pN}^{-1}$, $\lambda_2 = 0.08\ \text{pN}^{-1}$, $k_BT = 4.114$ pN·nm, $N = 5$, $m = 1$.

The sweeps below share a consistent picture: emergent catch behavior is governed by the competition between force-suppressed death and force-enhanced immigration, and is remarkably insensitive to standard rebinding. Catch ($d\tau_m/dF|_{F=0} > 0$) requires slow death (low $k^0_{\text{off}}$) together with strong force-driven

**Table S5.** Sweep ranges for phase diagrams.

| Parameter | Sweep range | Units |
|---|---|---|
| $k^0_{\text{off}}$ | [0.01, 1.0] | $s^{-1}$ |
| $k^0_{\text{on}}$ | [0.0, 0.5] | $s^{-1}$ |
| $\lambda_0$ | [0.0, 1.0] | $s^{-1}$ |
| $\lambda_1$ | [0.0, 0.3] | $pN^{-1}$ |
| $\lambda_2$ | [0.001, 1.0] | $pN^{-1}$ |
| $N$ | [2, 12] | — |

immigration (large $\lambda_1$ and small decay $\lambda_2$); the $k^0_{\text{off}}$–$\lambda_1$ and $k^0_{\text{off}}$–$\lambda_2$ boundaries are approximately hyperbolic, so a faster intrinsic off-rate can always be compensated by stronger or longer-lived immigration. The force-independent baseline $\lambda_0$ plays the same qualitative role as $\lambda_1$ and can sustain catch even at $\lambda_1 = 0$, but only at biologically large values ($\lambda_0 \gtrsim 0.5\ s^{-1}$). By contrast, every diagram involving the standard rebinding rate $k^0_{\text{on}}$ is essentially flat along that axis: with $\lambda_1 = 0.08\ s^{-1}\ pN^{-1}$ dominating the birth term, rebinding contributes negligibly and the catch/slip boundary is set by $k^0_{\text{off}}$, $\lambda_1$, and $\lambda_2$ alone. This mirrors the structural non-identifiability of $(k^0_{\text{on}}, \Delta x_{\text{on}})$ found in the Bayesian inference (Appendix S6): the same parameters that are statistically unconstrained by the data are also dynamically irrelevant to the phase boundary.

Multivalency systematically enlarges the catch region. Increasing $N$ shifts the $k^0_{\text{off}}$ boundary toward higher death rates, lowers the $\lambda_1$ threshold required for catch, and relaxes the upper bound on the immigration decay $\lambda_2$ — in each $N$-dependent panel, the catch domain grows monotonically with $N$. This is the signature of the cooperative, network-level mechanism: more residue-binding contributions provide more opportunities for force-driven re-engagement, so the collective bond tolerates faster individual ruptures while still producing a net lifetime increase with force. Taken together, the diagrams show that CFPR catch behavior is not a finely tuned corner of parameter space but a broad, robust regime accessible across physiologically plausible values, controlled by a small number of immigration and death parameters and largely decoupled from rebinding.

#### Phase diagrams: death–immigration coupling

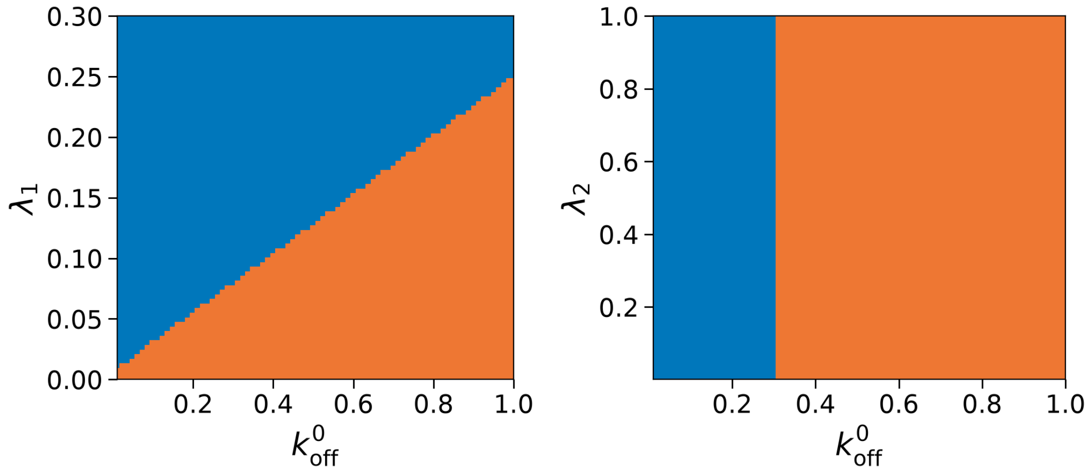


**Figure S13. Left:** $k^0_{off}$ vs. $\lambda_1$. Catch requires slow death (low $k^0_{off}$) and strong force-enhanced immigration (high $\lambda_1$); the boundary is approximately hyperbolic. **Right:** $k^0_{off}$ vs. $\lambda_2$. Catch persists when death is slow and immigration decays slowly (low $\lambda_2$). ■ catch regime (> 0), ■ slip regime (≤ 0).

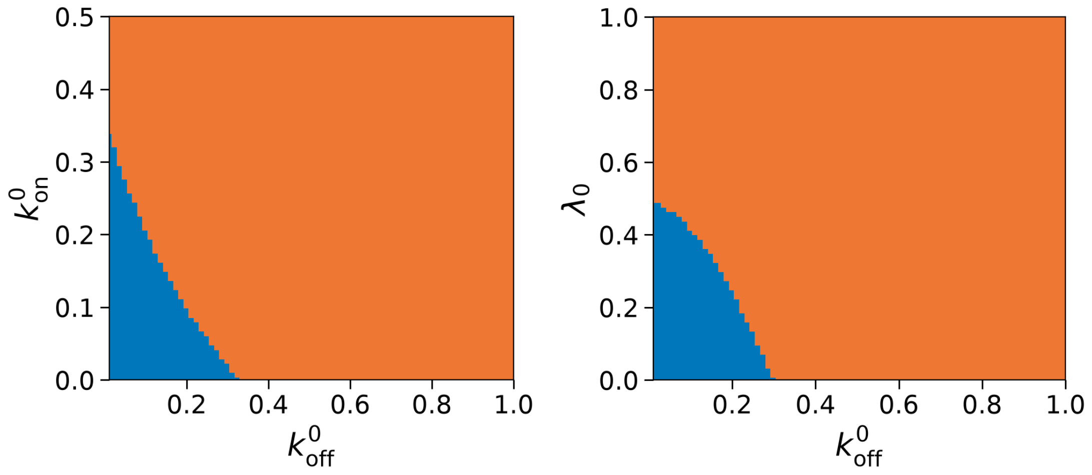


**Figure S14. Left:** $k^0_{off}$ vs. $k^0_{on}$. With $\lambda_1 = 0.08\ \mathrm{s}^{-1}\,\mathrm{pN}^{-1}$ dominating the birth rate, the catch boundary is insensitive to $k^0_{on}$ and controlled primarily by $k^0_{off}$. **Right:** $k^0_{off}$ vs. $\lambda_0$. A force-independent baseline immigration $\lambda_0$ extends the catch regime to higher death rates. ■ catch regime (> 0), ■ slip regime (≤ 0).

#### Phase diagrams: multivalency ($N$)

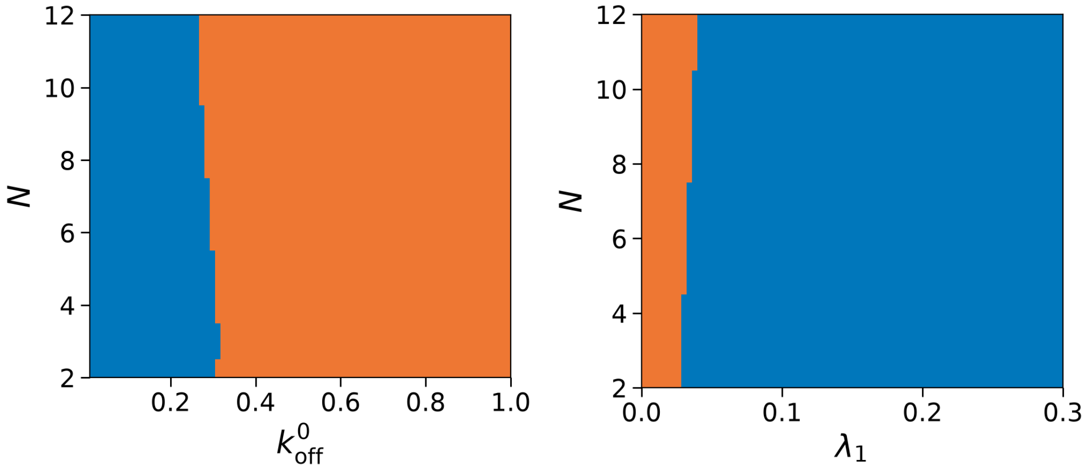


**Figure S15. Left:** $k^0_{\mathrm{off}}$ vs. $N$. Larger $N$ provides more rebinding opportunities, shifting the catch–slip boundary toward higher death rates. **Right:** $\lambda_1$ vs. $N$. Catch is accessible when either immigration is strong, or multivalency is high; the boundary is roughly hyperbolic in $(\lambda_1, N)$. ■ catch regime ($> 0$), ■ slip regime ($\leq 0$).

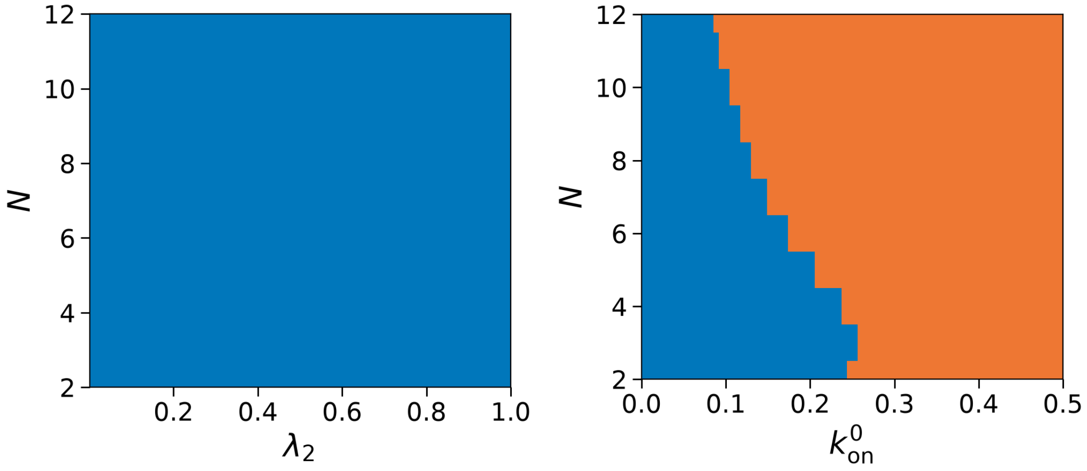


**Figure S16. Left:** $\lambda_2$ vs. $N$. The immigration decay rate $\lambda_2$ must be sufficiently small for catch to persist; larger $N$ relaxes this requirement. **Right:** $k^0_{\mathrm{on}}$ vs. $N$. With $\lambda_1 = 0.08\ \mathrm{s}^{-1}\ \mathrm{pN}^{-1}$, rebinding contributes weakly; the diagram is nearly uniformly catch across $k^0_{\mathrm{on}}$. ■ catch regime ($> 0$), ■ slip regime ($\leq 0$).

#### Phase diagrams: immigration parameters

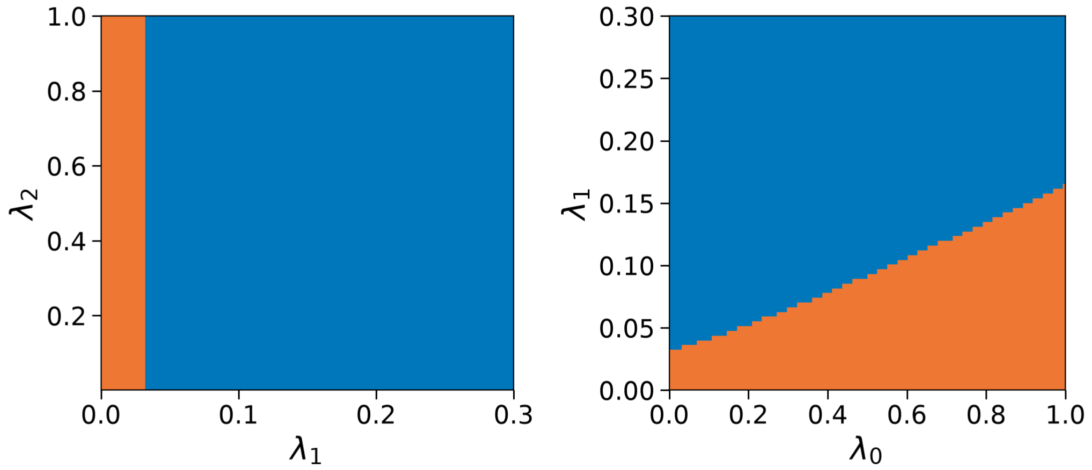


**Figure S17. Left:** $\lambda_1$ vs. $\lambda_2$. Catch requires $\lambda_1 > 0$ and $\lambda_2$ below a threshold ($\approx 0.2$ pN$^{-1}$); the boundary is nearly vertical. **Right:** $\lambda_0$ vs. $\lambda_1$. Force-independent immigration $\lambda_0$ can drive catch even at $\lambda_1 = 0$, but only at biologically large values ($\lambda_0 \gtrsim 0.5$ s$^{-1}$). ■ catch regime ($> 0$), ■ slip regime ($\leq 0$).

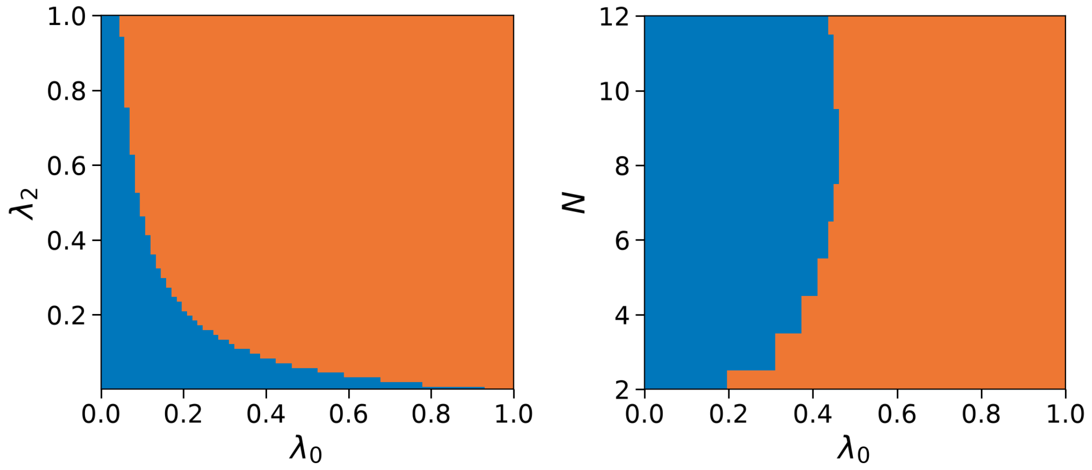


**Figure S18. Left:** $\lambda_0$ vs. $\lambda_2$. Catch requires either a large baseline immigration or a small decay rate $\lambda_2$. **Right:** $\lambda_0$ vs. $N$. Moderate $\lambda_0$ combined with large $N$ can sustain catch behavior. ■ catch regime ($>$ 0), ■ slip regime ($\leq 0$).

#### Phase diagrams: rebinding parameters

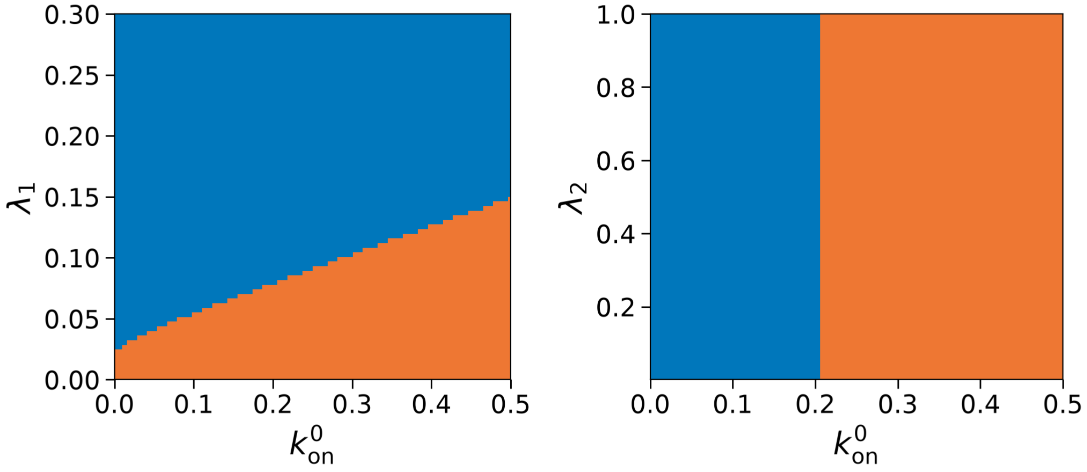


**Figure S19. Left:** $k_{on}^0$ vs. $\lambda_1$. The catch boundary is dominated by $\lambda_1$ and essentially independent of $k_{on}^0$. **Right:** $k_{on}^0$ vs. $\lambda_2$. Rebinding plays a minor role; the boundary is nearly vertical and is controlled by $\lambda_2$. ■ catch regime (> 0), ■ slip regime (≤ 0).

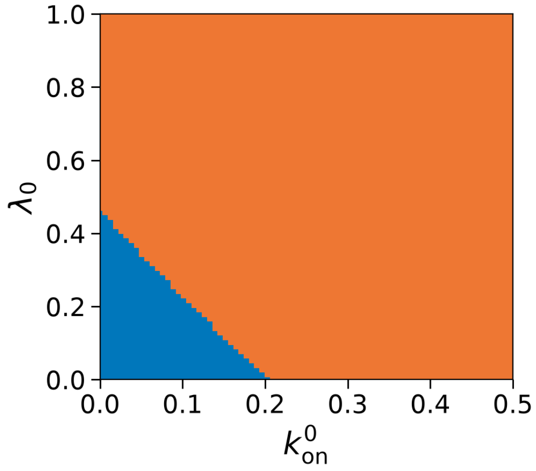


**Figure S20.** $k_{on}^0$ vs. $\lambda_0$. With $\lambda_1 = 0.08\ s^{-1}\ pN^{-1}$, the catch/slip boundary is essentially independent of both rebinding parameters. ■ catch regime (> 0), ■ slip regime (≤ 0).
